# Interferon gamma-induced MX1 and IFITM1 inhibit Varicella-Zoster Virus Replication

**DOI:** 10.64898/2026.09.28.754940

**Authors:** Seong K. Kim, Akhalesh K. Shakya, Hongyan Guo

**Affiliations:** Department of Microbiology and Immunology, and Center for Molecular and Tumor Virology, Louisiana State University Health-Shreveport, Shreveport, Louisiana 71130-3932

**Keywords:** varicella-zoster virus, interferon gamma, interferon response factor 1, JAK inhibitor, replication inhibition, IE62, MX1, IFITM1, keratinocyte

## Abstract

Interferon gamma (IFN-γ) is a potent cytokine secreted following primary varicella-zoster virus (VZV) infection. Our previous studies demonstrated that pretreatment with IFN-γ completely inhibited VZV replication in lung fibroblast MRC-5 and retinal epithelial ARPE-19 but not in melanoma MeWo cells, suggesting that IFN-γ-stimulated protein(s) inhibit viral replication. Our microarray analysis revealed that 34 interferon-stimulated genes (ISGs) were upregulated by greater than 2.5-fold at 8 h post-treatment in both ARPE-19 and MRC-5 cells compared to those of MeWo cells. The depletion of CTSS, GBP4, MX1, IFIT1, IFITM1 and IRF1 by siRNA in IFN-γ-treated cells significantly increased VZV yields. Ectopic overexpression of interferon-induced myxovirus resistance 1 (MX1) or transmembrane protein 1 (IFITM1) significantly reduced VZV yield both in ARPE-19 and MeWo cells. IFITM1 abrogated viral major immediate-early IE62 protein-mediated *trans*-activation. IFITM1 reduced the level of VZV immediate-early IE62 protein, but not the IE62 mRNA, suggesting that IFITM1 expression reduces the expression level of IE62 by post-transcriptional regulation. Both human primary epidermal keratinocytes and calcium chloride-differentiated keratinocytes exhibit high levels of endogenous MX1 gene expression. Pharmacologic inhibition of JAK signaling reduced MX1 expression and enhanced VZV replication in calcium chloride-induced differentiated keratinocytes. Collectively, these findings identify MX1 and IFITM1 as key IFN-γ-induced restriction factors that inhibit VZV replication and suggest that suppression of JAK-STAT signaling can promote VZV replication in differentiated keratinocytes.

**IMPORTANCE:** Varicella-zoster virus (VZV) is a ubiquitous human pathogen that causes chickenpox and shingles, which are characterized by the formation of fluid-filled skin lesions. Our results showed that MX1 and IFITM1 inhibited VZV replication both in ARPE-19 and MeWo cells. Unexpectedly, high levels of endogenous MX1 were detected in human primary epidermal Keratinocytes. Suppressing MX1 expression with a JAK inhibitor increased VZV replication in calcium chloride-treated keratinocytes. These results indicate that both the expression level of endogenous MX1 and the differentiation status of the keratinocytes influence VZV replication in keratinocytes. Understanding the mechanisms by which MX1 and IFITM1 play roles in VZV gene programming may be important in determining the tissue restriction of VZV.

## INTRODUCTION

Varicella-zoster virus (VZV) is a human alphaherpesvirus that causes varicella (also known as chickenpox) in primary lytic infection and herpes zoster (shingles) during reactivation from latent infection [1, 2]. The viral genome is ∼125 kbp in size and encodes at least 70 unique open reading frames (ORFs) [1]. The VZV immediate-early 62 protein (IE62) initiates VZV replication by transactivating viral immediate-early (IE), early (E), and late (L) genes [1, 3].

Insulin-degrading enzyme (IDE) is a receptor for VZV infection [4] and is a ubiquitous protein [5], which contributes to the broad tissue tropism of VZV. VZV induces signal transducer and activator of transcription 3 (STAT3) activation, which results in the expression of the antiapoptotic protein survivin as well as the inhibition of IFN-α and STAT1 [6]. VZV IE62 antagonizes type I IFN induction by inhibiting IFN regulatory factor 3 (IRF3) phosphorylation [7].

Interferon gamma (IFN-γ), the lone member of type II IFN [8], produced during viral infection stimulates transcription of cellular genes that mediate antiviral responses against several herpesviruses [9–11]. IFN-γ is primarily produced by Natural Killer (NK) cells and T lymphocytes (Th1 and CD8+ cells) during innate and adaptive immune responses, while its capacity to respond is widespread among almost all nucleated cells in the body [12, 13]. IFN-γ induces varying levels of antiviral interferon-stimulated genes (ISGs) in different cell types due to cell-specific receptor densities, baseline epigenetic states, variations in JAK/STAT signaling, and the presence of cell-intrinsic transcriptional regulators [14, 15]. IFN-γ is a potent cytokine produced following primary VZV infection [16 17] and inhibits VZV production in human neurons [8], human embryonic lung fibroblasts [18], and retinal epithelial ARPE-19 cells [19]. Furthermore, VZV reactivation correlates with a decline in IFN-γ-producing immune cells [20]. The IFN-γ receptor (IFN-γR) consists of ligand-binding IFN-γRα chains associated with Janus tyrosine kinase (JAK)1 and two signal-transducing IFN-γRβ chains associated with JAK2 [21, 22]. Binding of IFN-γ to its receptor activates JAK1 and JAK2. Activated JAK1 phosphorylates the IFN-γRα chain to create a docking site for STAT1 binding and phosphorylation. This phosphorylation leads to the translocation of STAT1 homodimers to the nucleus, where they bind to gamma-activated sequence (GAS) sites on the promoters of downstream target genes [21, 22]. One of the primary response genes transactivated by IFN-γ signaling is the transcription factor IFN response factor 1 (IRF1) which activates a large number of secondary response genes, known as ISGs [23, 24].

The interferon-induced protein with tetratricopeptide repeats (IFIT) gene family is interferon-stimulated genes (ISGs) for their antiviral properties [25]. The human IFIT gene family consists of four genes *IFIT1, IFIT2, IFIT3*, and *IFIT5* [21]. Humans also have three antiviral IFITMs: IFITM1, IFITM2, and IFITM3 [26]. IFITM (IFITM1, IFITM2, and IFITM3) expression inhibits viral entry (HIV, influenza A virus, West Nile virus, and dengue virus) into cells [27, 28]. Inhibition of viral entry is thought to occur via IFITM-mediated changes in the physical characteristics of the host cell membrane thereby inhibiting virus-cell membrane fusion [29]. IFITM expression reduces HIV-1 protein synthesis by preferentially excluding viral mRNA transcripts from translation and thereby restricts viral production [30].

MX proteins are well-characterized interferon (IFN)-induced dynamin-like GTPases that are present in all vertebrates and have antiviral activities against a broad range of viruses [31, 32]. Mouse MX1 inhibits a number of negative-strand RNA viruses, including IAV, Thogoto virus, Dhori virus, and Batken virus [33, 34]. Mouse MX1 also inhibits growth of the human herpes simplex virus 1 (HSV-1) and HSV-2 [32, 35]. Human MX1 (hMX1). which is ∼65% identical to mMX1 at the amino acid level, is solely cytoplasmic and inhibits viruses replicating in the cytoplasm such as LaCrosse virus (LACV), as well as viruses replicating in the nucleus, like IAV and HSV-1 [36–38].

In our previous study, we showed that pre-treatment with IFN-γ inhibited VZV gene expression and replication [19], suggesting that IFN-γ-stimulated protein(s) inhibit viral replication. Because IFN-γ completely inhibits VZV replication both in ARPE-19 and MRC-5 cells but only weakly in MeWo cells, this cell-line-dependent difference facilitates the selection of candidate antiviral IFN-γ-stimulated genes. We performed Affymetrix microarray analyses to identify IFN-γ-stimulated genes that inhibit VZV replication and characterized functional mechanisms of the selected ISGs.

## RESULTS

### IFN-_γ_-stimulated gene(s) that inhibits VZV replication

IFN-γ is a potent cytokine produced in response to primary VZV infection [16, 17]. Furthermore, VZV reactivation correlates with a decline of VZV-specific IFN-γ-producing immune cells [20]. Our published data showed that IFN-γ significantly inhibited VZV replication in ARPE-19, MRC-5, and cells but poorly inhibited VZV replication in melanoma MeWo cells [19], suggesting that critical antiviral ISG(s) are absent in MeWo. This difference in cell lines provides a tool to identify ISGs required for VZV restriction.

To identify IFN-γ-stimulated genes that inhibit VZV replication, microarray analyses were performed (see in Materials & Methods). ARPE-19, MRC-5, and MeWo cells were treated with 20 ng/ml of human IFN-γ. The treated cells were harvested at 8 h post-treatment and used in DNA microarray analysis using GeneChip human Genome U133 Plus 2.0 (Affymetrix, Santa Clara, CA). A complete list of genes altered by IFN-γ treatment is available in Table S1. Table 1 lists 48 interesting ISGs that are upregulated by IFN-γ treatment. Of these genes, 34 ISGs were upregulated by greater than 2.5-fold at 8 h post-treatment in both ARPE-19 and MRC-5 cells compared to those of MeWo cells (Table 1). Some ISGs including chemokine genes were also weakly upregulated at 8 h post-treatment in both ARPE-19 and MRC-5 cells compared to those of MeWo cells (Table 1). IRF1 transcripts were strongly upregulated in all three cell lines, indicating that MeWo cells can respond to IFN-γ stimulation, albeit not to the same extent as the other cell lines. This is consistent with our previous results, which showed that IFN-γ treatment increased IRF1 expression levels approximately eightfold in ARPE-19, MRC-5, and MeWo cells [19]. These results suggest that IRF1 does not directly inhibit VZV replication. We hypothesize that one or more of the upregulated antiviral ISGs are responsible for the inhibitory effect of IFN-γ.

**Table 1.** Interferon-stimulated genes upregulated by IFN-γ at 8 h post-treatment.

| **Gene name** | Fold change<sup>a</sup> |  |  | Description |
|---|---|---|---|---|
|  | ARPE-19 | MRC-5 | MeWo |  |
| **APOL2** | 13.9 | 11.2 | 3 | apolipoprotein L, 2 |
| **BST2** | 28.8 | 23.4 | 5.3 | Bone marrow stromal cell antigen 2 |
| **CCL2** | 30.9 | 11.4 | 1.1 | chemokine (C-C motif) ligand 2 |
| **CIITA** | 20.8 | 7.6 | 1 | class II, major histocompatibility complex, transactivator |
| **CLIC2** | 7.4 | 15.2 | 1.4 | chloride intracellular channel 2 |
| **CMPK2** | 183.5 | 35.5 | 1.8 | cytidine monophosphate (UMP-CMP) kinase 2, mitochondrial |
| **CTSS** | 18.8 | 6.8 | 1.1 | cathepsin S |
| **CXCL9** | 0.9 | 67.2 | 1.2 | chemokine (C-X-C motif) ligand 9 |
| **CXCL10** | 4.5 | 369.6 | 0.8 | chemokine (C-X-C motif) ligand 10 |
| **CXCL11** | 2.9 | 256 | 0.9 | chemokine (C-X-C motif) ligand 11 |
| **DDX58** | 30 | 3.7 | 1.5 | DEAD (Asp-Glu-Ala-Asp) box polypeptide 58 |
| **DDX60** | 13.9 | 3.3 | 1 | DEAD (Asp-Glu-Ala-Asp) box polypeptide 60 |
| **EPSTI1** | 29.7 | 25.5 | 1.1 | epithelial stromal interaction 1 (breast) |
| **FBXO6** | 14.6 | 5.1 | 0.9 | F-box protein 6 |
| **GBP1** | 19.4 | 13.5 | 1.4 | guanylate binding protein 1 |
| **GBP2** | 64 | 41.6 | 16.2 | guanylate binding protein 2 |
| **GBP4** | 22.6 | 22.8 | 1.2 | guanylate binding protein 4 |
| **GBP5** | 17.8 | 10.9 | 1 | guanylate binding protein 5 |
| **IFI27** | 27.1 | 6.2 | 1.1 | interferon, gamma-inducible protein 27 |
| **IFI30** | 29.9 | 23.6 | 2.7 | interferon, gamma-inducible 30 |
| **IFI35** | 20.3 | 11.2 | 2.9 | interferon, gamma-inducible 3=5 |
| **IFI44L** | 24.2 | 48.3 | 1.3 | interferon-induced protein 44 like |
| **IFIT1** | 33.8 | 11.2 | 2.2 | interferon-induced protein with tetratricopeptide repeats 1 |
| **IFIT2** | 81 | 26.5 | 4.2 | interferon-induced protein with tetratricopeptide repeats 2 |
| **IFIT3** | 42.8 | 18.6 | 6.6 | interferon-induced protein with tetratricopeptide repeats 3 |
| **IFIT5** | 4.2 | 3.4 | 2.2 | interferon-induced protein with tetratricopeptide repeats 5 |
| **IFITM1** | 44.6 | 6 | 2 | interferon induced transmembrane protein 1 |
| **IFITM2** | 2.7 | 1.6 | 1.4 | Interferon induced transmembrane protein 2 |
| **IFITM3** | 2.5 | 1.5 | 1.2 | Interferon induced transmembrane protein 3 |
| **IGFBP5** | 22.5 | 1.3 | 1.2 | insulin like growth factor binding protein 5 |
| **IRF1** | 63.1 | 28.6 | 33.1 | interferon response factor 1 |
| **IRF7** | 14.1 | 2.3 | 2.8 | interferon regulatory factor 7 |
| **ISG15** | 32.2 | 4.5 | 2.5 | interferon stimulated protein (=ubiquitin-like modifier) |
| **ISG20** | 38.5 | 5.5 | 1.3 | interferon stimulated exonuclease protein |
| **MX1** | 192.7 | 38 | 1 | Myxovirus (influenza virus) resistance 1 |
| **NAMPT** | 18.6 | 6.9 | 1.2 | nicotinamide phosphoribosyltransferase |
| **OAS1** | 71.6 | 2.6 | 2.6 | 2'-5' oligoadenylate synthetase 1 |
| **OAS2** | 37.4 | 4.1 | **1** | 2'-5' oligoadenylate synthetase 2 |
| **OASL** | 12.4 | 1.7 | 1 | 2'-5'-oligoadenylate synthetase-like |
| **RARRES3** | 60.1 | 56.9 | 3.2 | retinoic acid receptor responder (tazarotene induced) 3 |
| **RSAD2** | 182.3 | 1.5 | 1 | radical S-adenosyl methionine domain containing 2 |
| **RSPO3** | 18.6 | 6.9 | 1.2 | R-spondin 3 |
| **SAMHD1** | 21.1 | 10.2 | 1.5 | SAM domain and HD domain 1 |
| **SERPING1** | 9.8 | 21.3 | 1.1 | serpin peptidase inhibitor, clade G (C1 inhibitor), member 1 |
| **TNFSF10** | 26.9 | 10.7 | 1.1 | tumor necrosis factor (ligand) superfamily, member 10 |
| **TNFSF13B** | 5.7 | 11.6 | 2.6 | tumor necrosis factor (ligand) superfamily, member 13b |
| **WARS** | 11.9 | 6.7 | 13.7 | tryptophanyl-tRNA synthetase |
| **XAF1** | 63.6 | 9.2 | 2.1 | XIAP associated factor 1 |

**a**
Presented as the mean fold change from three replicate experiments at 8 hpt. A549, ARPE-19, and MeWo cells were treated with 0 or 20 ng/mL of human IFN-γ (Cell Sciences, MA), harvested at 8 h post-treatment, and used in DNA microarray analyses using GeneChip human Genome U133 Plus 2.0 (Affymetrix, Santa Clara, CA). The mean fold change was calculated as the ratios of the average gene expression levels between the treated and untreated samples.

### Depletion of CTSS, GBP4, MX1, IFIT1, and IFITM1 in IFN-_γ_-treated cells increases VZV replication

Our previous results showed that the knockdown of IRF1 in IFN-γ-treated ARPE-19 cells leads to an increase in VZV replication [19]. To identify IFN-γ-stimulated gene(s) that inhibit VZV replication, small interfering RNA (siRNA) knockdown of the upregulated ISGs in ARPE-19 cells was performed in the context of VZV infection. To deplete antiviral gene expression, ARPE-19 cells were reverse transfected with siGenome SMARTpool siRNAs targeting 11 ISGs (CTSS, GBP4, IFIT1, IFITM1, IRF1, IRF7, ISG15, MX1, OAS2, RSAD2, SAMHD1; G-CUSTOM-430650 and 433679, Dharmacon, CO) or a nontargeting control (siNTC). The cells were treated with 20 ng/ml of IFN-γ at 8h post-transfection (hpt) and harvested at 24 h post-treatment (hpt) for western blot analyses. Knockdown efficiencies of 11 ISGs were 57 to 93% in western blot analysis (Fig. S1). Depletion of IRF1, CTSS, GBP4, MX1, IFIT1, and IFITM1 in IFN-γ-treated cells significantly increased VZV yields (Fig. 1A).

**FIG 1.**
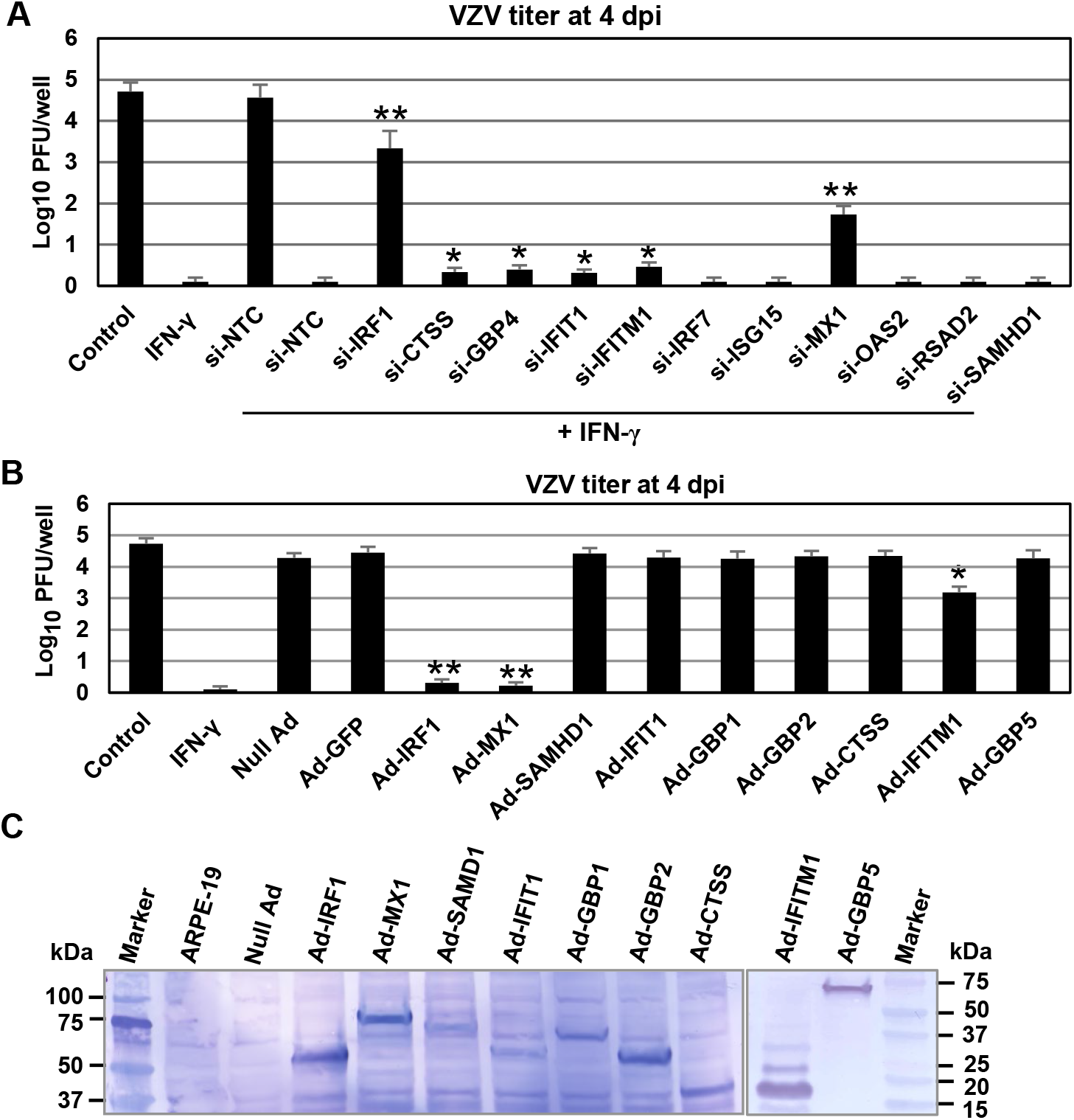
MX1 and IFITM1 reduce VZV yield in ARPE-19 cells. **(A)** Depletion of CTSS, GBP4, MX1, IFIT1, and IFITM1 in IFN-γ-treated cells increases VZV replication. ARPE-19 cells (3.5×10^5^, 12-well plates) were transfected with siRNA targeting 11 ISGs (G-CUSTOM-430650, Dharmacon, CO) or a nontargeting control (si-NTC, Dharmacon). At 8 hours post-transfection, the cells were treated with 20 ng/ml of IFN-γ and were infected with 0.01 MOI of VZV (AV92-3:L, ATCC) at 24 hours post-treatment (hpt). Cell-associated infectious VZV titers were determined by plaque assay at 4 dpi. Data are the averages of three independent experiments. Control: untreated ARPE-19 cells. *, P<0.05 for comparison with the si-NTC. **, P<0.01 for comparison with the si-NTC. **(B)** ARPE-19 cells were infected with 30 MOI of Adenovirus expressing ISGs. The cells were infected with 0.01 MOI of cell-free VZV 24 hours later. Virus titers were determined by plaque assay at 4 days post-infection (dpi). Control, untreated ARPE-19 cells; IFN-γ, IFN-γ-treated cells. **, P<0.01 for comparison with the null Ad. **(C)** Western blot analysis of ISGs. ARPE-19 cells were infected with 30 MOI of recombinant adenoviruses expressing ISGs with a C-terminal Flag tag (Null Ad, Ad-GFP, Ad-IRF1, Ad-MX1, Ad-SAMHD1, Ad-IFIT1, Ad-GBP1, Ad-GBP2, Ad-CTSS, Ad-GBP5, and Ad-IFITM1), and the cells were harvested at 48 h post-infection for western blot analyses using anti-flag antibody. All of the proteins (IRF1 [48 kDa], MX1 [75 kDa], SAMHD1 [72 kDa], IFIT1 [56 kDa], GBP1 [68 kDa], GBP2 [65 kDa], CTSS [37 kDa], IFITM1 [17 kDa], and GBP5 [67 kDa]) were expressed at their expected sizes. The sizes of molecular mass standards (in kilodaltons) (Prestained protein ladders; Bio-Rad Laboratories, CA) are shown.

One of the major primary response genes transactivated by IFN-γ–activated JAK/STAT signaling is the transcription factor IFN response factor 1 (IRF1) that in turn activates many secondary response genes including antiviral ISGs [23, 24]. We showed that infection with adenovirus expressing IRF1 (Ad-IRF1) completely inhibited VZV replication in ARPE-19 cells [19]. However, infection with null Ad or Ad-SAMHD1 reduced virus yields modestly [19]. To investigate whether the upregulated ISGs reduce VZV yield, ARPE-19 cells were infected with the ISG-expressing recombinant adenoviruses (Ad-IRF1, Ad-MX1, Ad-SAMHD1, Ad-IFIT1, Ad-GBP1, Ad-GBP2, Ad-CTSS, Ad-GBP5, and Ad-IFITM1 and subsequently infected with 0.01 MOI of cell-free VZV at 24 hpi. At 4 days post-infection (dpi), intracellular VZV was measured by plaque assay. All expressed ISGs were expected size (Fig. 1C). MX1 as well as IRF1 completely reduced intracellular VZV yields (Fig. 1B). IFITM1 reduced by 13-fold intracellular VZV yields compared to that of null Ad (Fig. 1B). However, the other six adenovirus expressing ISGs did not reduce virus yields.

### IFITM1 inhibits VZV replication both in ARPE-19 and MeWo cells

To investigate whether ectopic expression of IFITM1 results in inhibition of VZV replication in MeWo cells, ARPE-19 and MeWo cells were treated with IFN-γ, infected with Ad-IRF1 or Ad-IFITM1, and subsequently infected with 0.01 MOI of cell-free VZV at 24 h post-IFN-γ treatment or 48 h post adenovirus-infection. Ectopic overexpression of IFITM1 reduced intracellular VZV yield in ARPE-19 and MeWo cells by 12- and 3-fold, respectively, compared with control null Ad (Fig. 2A and 2B, respectively). Ectopic overexpression of IRF1 almost completely reduced VZV yield in ARPE-19 cells, but only weakly reduced it in MeWo cells compared with the control null Ad (Fig. 2A and 2B, respectively), a finding consistent with our previously published results [19]. Western blot assay with anti-IRF1 antibody detected a protein corresponding to the size of flag-tagged IRF1 in the cells (Fig. 2C and 2D), confirming that IRF1-flag was expressed at high levels both in ARPE-19 and MeWo cells. IFITM1-flag was expressed at high levels in Ad-IFITM1-infected ARPE-19 cells, but only weakly in MeWo cells (Fig. 2C and 2D).

**FIG 2.**
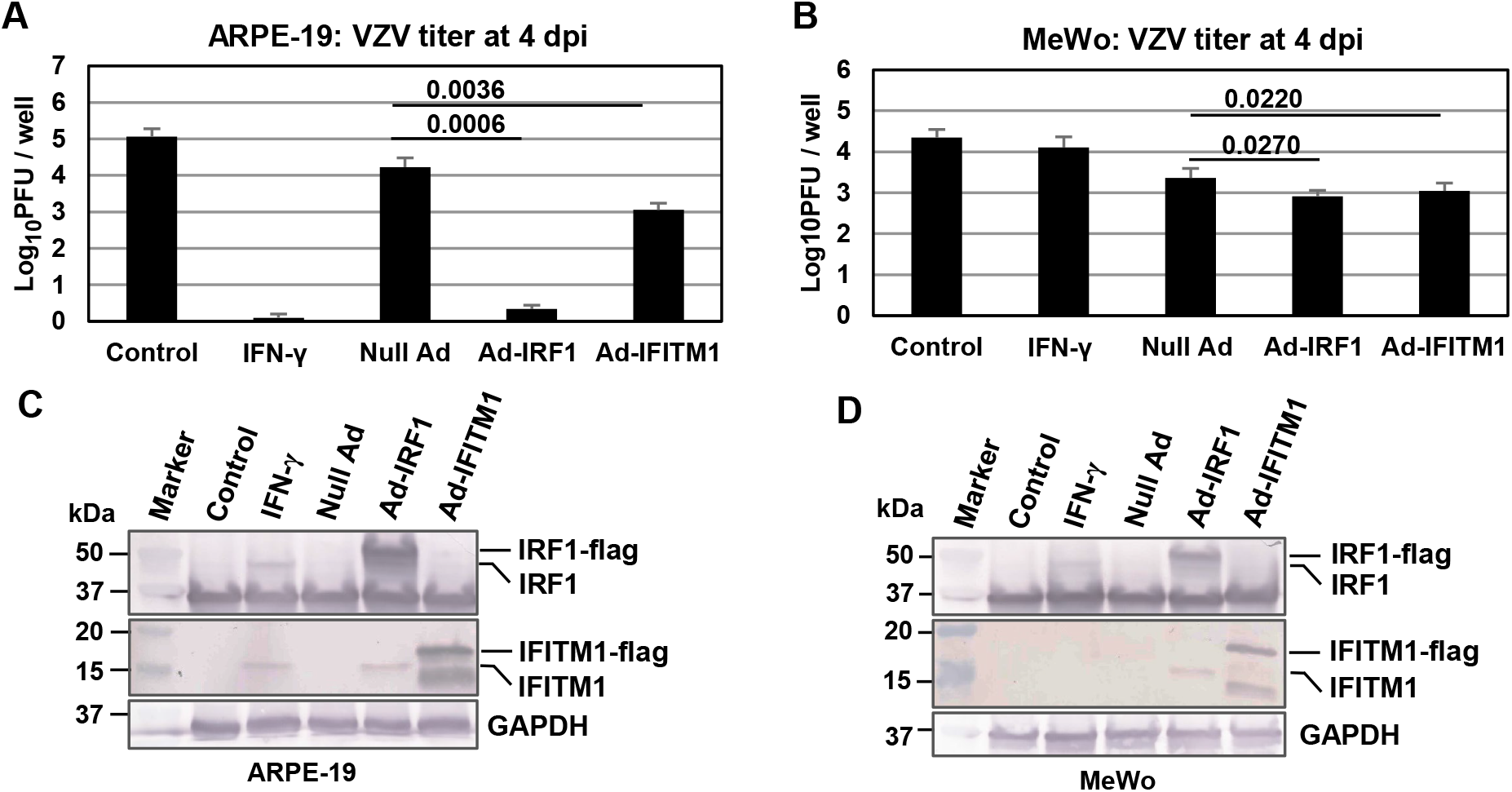
IFITM1 weakly inhibits VZV replication. **(A to D)** ARPE-19 and MeWo cells were plated in 24-well plates 3.5×10^5^), treated with 20 ng/ml of IFN-γ or infected with 30 MOI of recombinant adenoviruses (null Ad, Ad-IRF1, or Ad-IFITM1). The cells were infected with cell-free VZV at an MOI of 0.01 at 24 h post IFN-γ-treatment or 48 h post adenovirus-infection. Virus titers were determined by plaque assay at 4 dpi (A and B). Control, untreated ARPE-19 or MeWo cells. P values are the result of Student’s t tests and error bars indicate the standard deviation from three independent experiments. At 24 post IFN-γ-treatment or 48 h post Ad infection, the cells were harvested and used for western blot analyses with antibodies to IRF1, IFITM1, and GAPDH. (C and D).

### IRF1 and IFITM1 abrogate IE62-mediated *trans*-activation

Our previous results showed that IFN-γ inhibits VZV replication as well as the *trans*-activation activity mediated by VZV immediate-early protein (IE62) [19]. We also showed that adenovirus expressing IRF1 (Ad-IRF1) inhibited VZV replication by 4,000-fold in ARPE-19 cells, but reduced virus yields by only 3-fold in MeWo cells [19]. Luciferase reporter assays were performed with VZV promoter reporter plasmids and Ad-IRF1 (Fig. 3A and 3B). As expected, IFN-γ abrogated VZV IE62’s *trans*-activation activity of VZV ORF61 promoter by 98% in ARPE-19 (Fig. 3A, bar 4), but only by 34% in MeWo cells (Fig. 3B, bar 4). IE62’s *trans*-activation activity was reduced by 95% in ARPE-19 cells infected with Ad-IRF1 as compared to cells infected with control Ad-GFP (Fig. 3A, bar 10). Ectopic expression of IRF1 abrogated IE62’s *trans*-activation activity by 32% in MeWo cells (Fig. 3B, bar 10). The expression levels of IRF1 and GFP proteins were examined by western blot, and similar expression levels of IRF1 and GFP proteins were observed in each cell type (Fig. 3C and 3D). These results demonstrated that IRF1 inhibits IE62-mediated *trans*-activation in a cell line-dependent manner.

**FIG 3.**
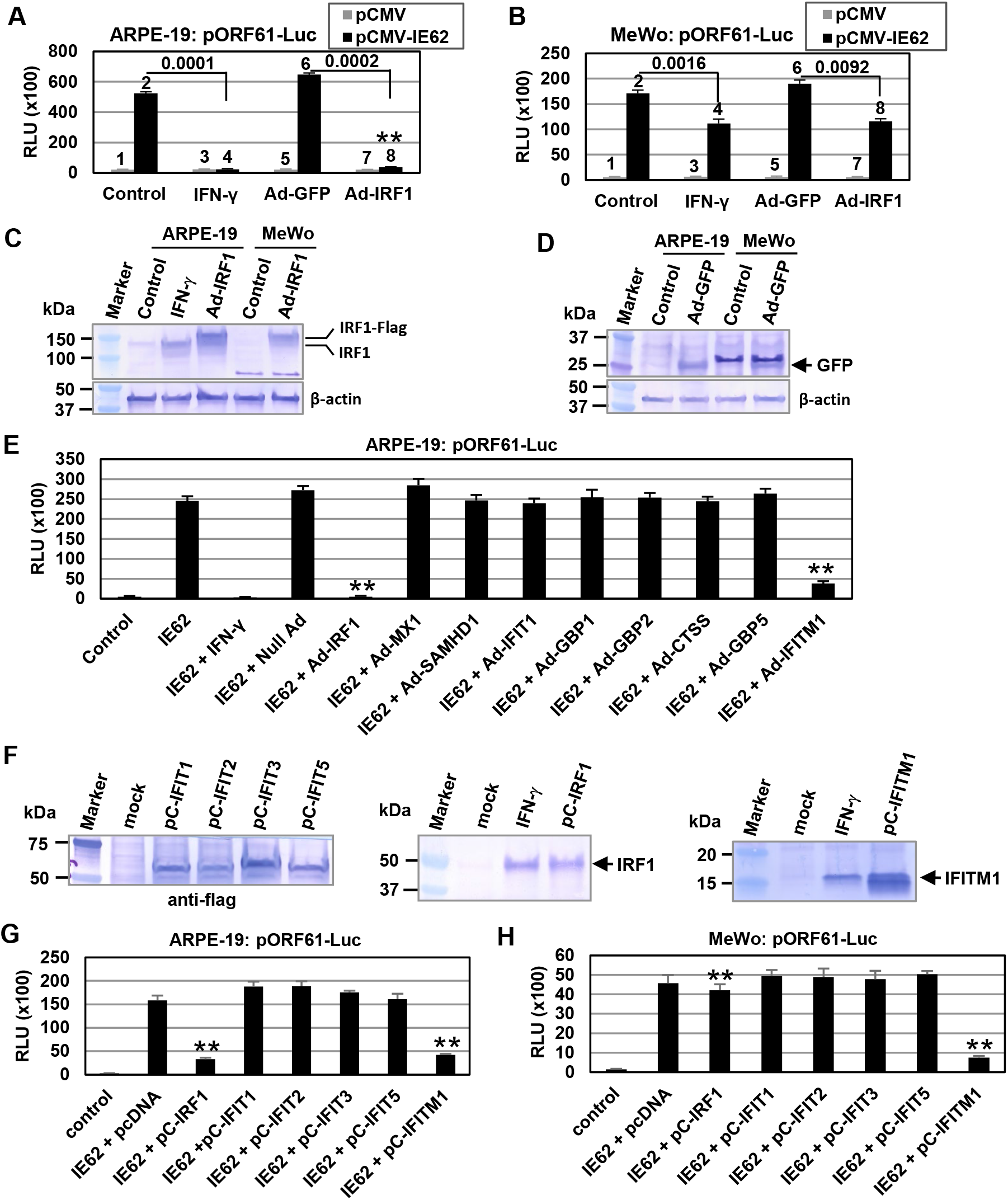
IRF1 and IFITM1 abrogate IE62-mediated *trans*-activation. ARPE-19 **(A)** and MeWo **(B)** cells were treated with 20 ng/ml of IFN-γ or infected with 10 MOI of recombinant adenoviruses (Ad-GFP or Ad-IRF1). At 24 h post IFN-γ-treatment or 48 h post adenovirus-infection, the cells were cotransfected with 0.07 pmol of VZV reporter plasmids pORF61-Luc and 0.07 pmol of effector plasmids (control empty vector pCMV or pCMV-IE62) at 24 hpi. At 40 h post-transfection, luciferase activity was measured with a luciferase assay kit (Promega, Madison, WI) and a Polarstar Optima plate reader. RLU, relative luminescence units. Reporter plasmid pICP22-SEAP was used to assess transfection efficiency. The firefly luciferase signals were normalized to the internal secreted alkaline phosphatase (SEAP) transfection control. *, P values are the result of Student’s t tests and error bars indicate the standard deviation from three independent experiments. Bars were labeled with a number on top. **(C and D)** Detection of IRF1 and GFP expression in ARPE-19 and MeWo cells. ARPE-19 and MeWo cells were treated with 20 ng/ml of IFN-γ or infected with 10 MOI of recombinant adenoviruses (Ad-GFP or Ad-IRF1). At 24 h post IFN-γ-treatment or 48 h post adenovirus-infection, the cells were harvested for western blot analyses using anti-IRF1, anti-GFP, and anti-GAPDH. **(E)** IFITM1 abrogates IE62-mediated *trans*-activation in ARPE-19 cells. ARPE-19 cells were treated with 20 ng/ml of IFN-γ or infected with 10 MOI of Adenovirus expressing ISGs. Twenty-four hours later, the cells were cotransfected with 0.07 pmol of VZV reporter plasmids pORF61-Luc and 0.07 pmol of effector plasmids (control empty vector pcDNA or pCMV-IE62). At 40 h post-transfection, luciferase activity was measured with a luciferase assay kit (Promega, Madison, WI). The firefly luciferase signals were normalized to the internal secreted alkaline phosphatase (SEAP) transfection control. Data are the averages of three independent experiments. **, P<0.01 for comparison with the null Ad. **(F)** Detection of IFIT proteins with an N-terminal 3XFlag tag. ARPE-19 cells were transfected with 0.2 pmol of pC-FIT1, pC-FIT2, pC-FIT3, pC-FIT5, pC-IFITM1 (Addgene, Warertown, MA; [77]), pC-IRF1 (pCMV6-XL5, Origene Tech., Rockville, MD), and the cells were harvested at 48 h post-transfection for western blot analyses using anti-flag (Cell Signaling, Inc., MA), anti-IFITM1 (Proteintech, IL), anti-IRF1 antibodies. **(G and H)** IFITM1 abrogates IE62-mediated *trans*-activation both in ARPE-19 and MeWo cells. ARPE-19 and MeWo cells were cotransfected with 0.07 pmol of VZV reporter plasmids pORF61-Luc and 0.07 pmol of effector plasmids (pC-IFIT expression vectors and pCMV-IE62). At 40 h post-transfection, luciferase activity was measured with a luciferase assay kit (Promega, Madison, WI). The firefly luciferase signals were normalized to the internal secreted alkaline phosphatase (SEAP) transfection control. control or pcDNA, pcDNAI/AMP; IE62, pCMV-IE62. Data are the averages of three independent experiments. **, P<0.01 for comparison with the pcDNA.

Luciferase reporter assays were carried out in ARPE-19 cells with the eight recombinant adenoviruses. Interestingly, IRF1 as well as IFITM1 significantly abrogated VZV IE62’s *trans*-activation activity of the immediate-early ORF61 promoter in ARPE-19 (Fig. 3E), but MX1 did not. The pORF61-Luc reporter was not transactivated by any of the nine recombinant adenoviruses expressing ISGs in the absence of IE62 (Fig. S2). IFIT proteins are supposed to confer immunity against viral infection [39]. We employed Five IFIT proteins IFIT1, IFIT2, IFIT3, IFIT5, and IFITM1 for luciferase reporter assays. IFITM2 and IFITM3, which were very weakly induced by IFN-γ, were not used for these experiments. Five IFIT proteins were expressed at high levels in ARPE-19 cells transfected with IFIT expression vectors containing the human cytomegalovirus (CMV) major immediate-early gene enhancer/promoter region (Fig. 3F). IRF1 significantly abrogated VZV IE62 *trans*-activation activity of the immediate-early ORF61 promoter in ARPE-19 cells (Fig. 3G), but not in MeWo cells (Fig. 3H). Interestingly, ectopic expression of only IFITM1 significantly abrogated VZV IE62 *trans*-activation activity both in ARPE-19 (Fig. 3G) and MeWo cells (Fig. 3H). In the absence of IE62, none of the five CMV IFIT effectors transactivated the pORF61-Luc reporter in ARPE-19 (Fig. S2B) or MeWo cells (Fig. S2C). These results suggest that IRF1 induce ISGs, such as IFITM1, that are involved in IE62-mediated trans-activation in ARPE-19 cells.

### IFITM1 reduces the expression level of IE62

To investigate the mechanism by which IFITM1 inhibits IE62 *trans*-activation activity and VZV replication, ARPE-19 cells were co-transfected with VZV IE62 expression plasmid pCIE62 and IFIT expression plasmids. The transfected cells were harvested at 48 hpt for western blot analyses using anti-IE62 antibody. Ectopic expression of IRF1 or IFITM1 significantly reduced IE62 expression levels to 33% and 37%, respectively (Fig. 4A; Quantification was done with Image Studio Lite software [LI-COR Biosciences, Lincoln, NE]). However, the other IFIT proteins IFIT1, IFIT2, IFIT3, and IFIT5 failed to reduce IE62 expression levels (Fig. 4A). To confirm these results, ARPE-19 and MeWo cells were co-transfected with pCIE62 and IFITM1. Interestingly, ectopic expression of IFITM1 significantly reduced IE62 expression to 23% in ARPE-19 cells (Fig. 4B) and 7% in MeWo cells (Fig. 4C). However, IRF1 did not reduce the expression level of IE62 in MeWo cells (Fig. 4C). These results indicated that IFITM1 inhibits VZV replication by reducing IE62 expression level.

**FIG 4.**
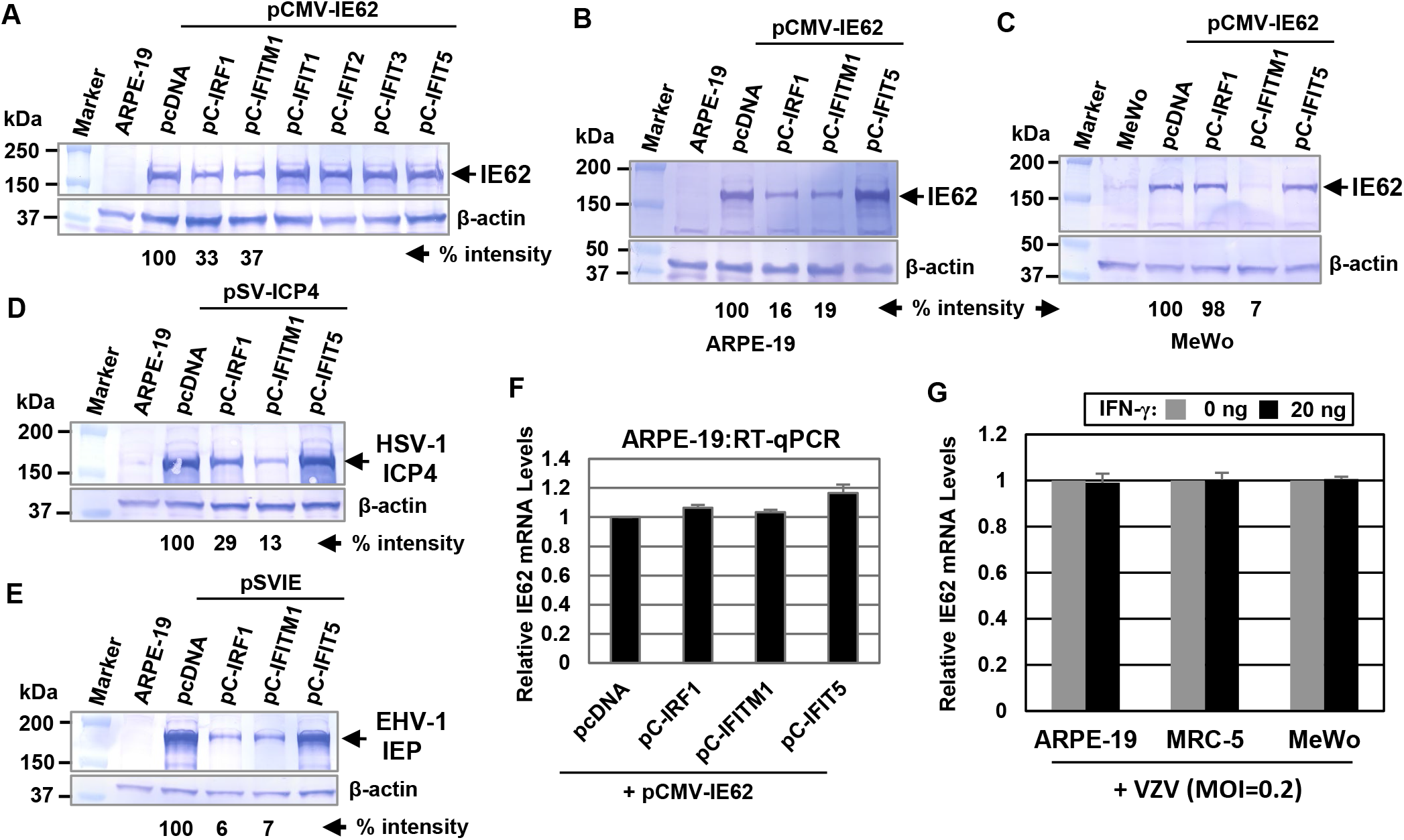
IFITM1 reduces VZV IE62 protein levels but does not affect IE62 mRNA levels in ARPE-19 cells. **(A)** ARPE-19 cells were co-transfected with 0.2 pmol of VZV IE62 expression plasmid pCIE62 and 0.3 pmol of IFIT expression plasmids and harvested at 48 h post-transfection for western blot analyses using anti-IE62 and β-actin antibodies. (**B and C**) IFITM1 reduces the expression levels of VZV IE62 in both ARPE-19 and MeWo cells. ARPE-19 and MeWo cells were co-transfected with 0.2 pmol of pCMV-IE62 and 0.3 pmol of pC-IRF1, pC-FITM1, or pC-FIT5, and the cells were harvested at 48 h post-transfection for western blot analyses. The densitometry analysis was performed with ImageJ (Rasband, W.S., NIH, Bethesda, Maryland). The intensities of the ISG bands were normalized to β-actin. (**D and E**) IFITM1 reduces the expression levels of herpes simplex virus 1 (HSV-1) ICP4 and equine herpesvirus 1 (EHV-1) IEP in ARPE-19 cells. ARPE-19 cells were co-transfected with 0.2 pmol of pSV-ICP4 or pSVIE and 0.3 pmol of pC-IRF1, pC-FITM1, or pC-FIT5, and the cells were harvested at 48 h post-transfection for western blot analyses using anti-ICP4 (58S, ATCC) and anti-IEP (OC33) antibodies. (**F**) ARPE-19 cells were co-transfected 0.2 pmol of pCMV-IE62 and 0.3 pmol of pC-IRF1, pC-FITM1, or pC-FIT5. Level of IE62 mRNA was measured by RT-qPCR at 48 h post-transfection. The data were normalized to GAPDH expression. Data are the averages of three independent experiments. (**G**) ARPE-19, MRC-5, and MeWo cells were treated with 0 or 20 ng of IFN-γ and infected with 0.2 MOI of cell-free VZV at 24 h post-transfection. IE62 mRNA levels were measured *by* RT-qPCR at 3 hpi. Data are the averages of three independent experiments. *, P<0.05 for comparison with the untreated control.

The alphaherpesvirus VZV IE62, herpes simplex virus 1 (HSV-1) immediate-early ICP4, and equine herpesvirus 1 (EHV-1) immediate-early IEP exhibit extensive homology [40–42]. When pC-IFITM1 was co-transfected with HSV-1 ICP4 expression vector pSV-ICP4 [43] or EHV-1 IEP expression vector pSVIE [44] in ARPE-19 cells, IFITM1 also significantly reduced the expression levels of ICP4 to 13% (Fig. 4D) and IEP to 7% (Fig. 4E).

### IFITM1 expression does not reduces the IE62 mRNA levels

VZV immediate-early protein IE62 was detected at 2 h; at both 2 h and 4 h, it appeared as distinct nuclear punctae located predominantly along the inner nuclear rim [3]. To measure the relative mRNA expression of IE62, ARPE-19 cells were co-transfected with pCIE62 and IFITM1. Level of IE62 mRNA was measured by quantitative real time PCR (RT-qPCR) at 48 hpt. It was interesting that ectopic expression of IRF1 as well as IFITM1 did not reduce the expression level of IE62 mRNA (Fig. 4F). ARPE-19, MRC-5, and MeWo cells were treated with 0 or 20 ng of IFN-γ and infected with 0.01 or 0.2 MOI of cell-free VZV at 24 hpt. IE62 mRNA levels were measured *by* RT-qPCR at 3 hpi. IFN-γ treatment did not reduce the expression level of IE62 mRNA (Fig. 4G and Fig. S2).

### Ectopic overexpression of human MX1 (MX1) inhibits VZV replication both in ARPE-19 and MeWo cells

To investigate whether ectopic expression of MX1 results in inhibition of VZV replication in MeWo cells, ARPE-19 and MeWo cells were treated with IFN-γ, infected with Ad-IRF1 or Ad-MX1, and subsequently infected with 0.01 MOI of cell-free VZV at 24 h post-IFN-γ treatment or 48 hpi. IFN-γ and Ad-IRF1 strongly induced MX1 expression in ARPE-19 cells (Fig. 5C), but not in MeWo cells (Fig. 5D). As expected, IFN-γ and Ad-IRF1 completely reduced intracellular VZV yields in ARPE-19 cells (Fig. 5A). However, IFN-γ and Ad-IRF1 reduced virus yields by only 3-fold in MeWo cells compared to control null Ad (Fig. 5B). Interestingly, Ad-MX1 infection completely reduced VZV yields in both ARPE-19 and MeWo cells (Figs. 5A and 4B). These results demonstrate that IFN-γ (and IRF1) weakly induces anti-VZV genes including MX1 in MeWo cells. Very similar results were obtained with the same alphaherpesvirus equine herpesvirus 1 (EHV-1). Ectopic overexpression of IRF1 strongly reduced EHV-1 yield in ARPE-19 cells but weakly reduced it in MeWo cells compared with the control null Ad (Fig. S3A and S3B, respectively). As expected, Ad-MX1 infection strongly reduced EHV-1 yields in both ARPE-19 and MeWo cells (Fig. S4), confirming that IFN-γ weakly induces anti-viral genes in MeWo cells. Ad-MX1 infection reduced VZV yields in a dose-dependent manner and almost completely inhibited VZV replication at an MOI of 30 in ARPE-19 cells (Fig. 5E, bars 5 and 6). As expected, IFITM1 weakly reduced VZV yield in ARPE-19 cells (Fig. 5E, bars 5 and 6). MX1 synergistically inhibited VZV replication with IFITM1 in ARPE-19 cells (Fig. 5E, bars 11 and 12).

**FIG 5.**
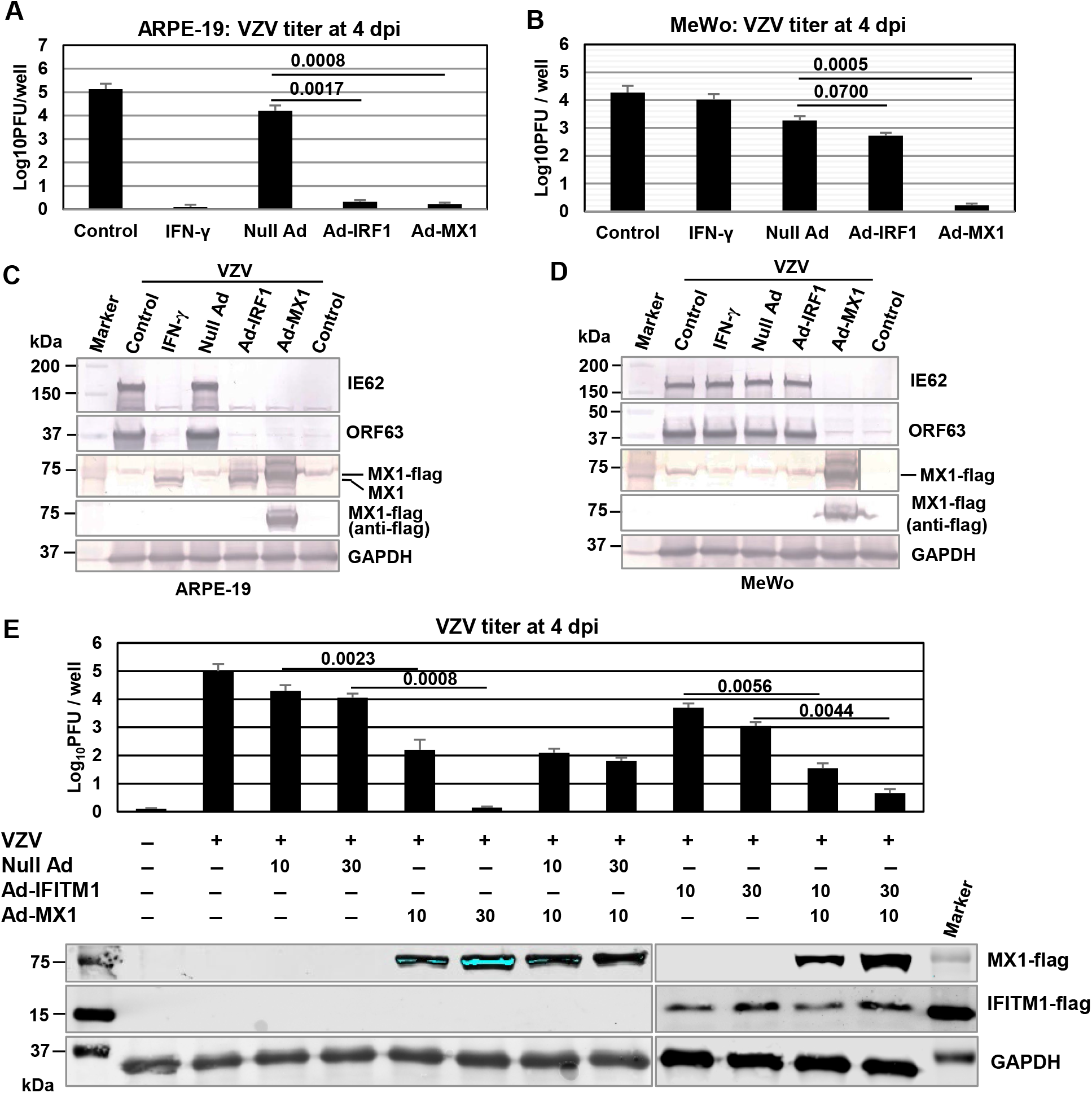
MX1 completely inhibits VZV replication both in ARPE-19 and MeWo cells. **(A and B)** ARPE-19 and MeWo cells were plated in 24-well plates (3.5×10^5^), treated with 20 ng/ml of IFN-γ or infected with 30 MOI of recombinant adenoviruses (null Ad, Ad-IRF1, or Ad-MX1) as described in Figure 1 and the cells were infected with cell-free VZV at an MOI of 0.01 at 24 h post IFN-γ-treatment (or 48 h post adenovirus-infection). Virus titers were determined by plaque assay at 4 dpi. Control, untreated ARPE-19 cells. **(C and D)** At 24 hpt (or 48 hpi), the cells were harvested for western blot analyses with antibodies to VZV IE62, VZV ORF63, MX1, and Flag. The sizes of molecular mass standards (in kilodaltons) (Bio-Rad Laboratories, CA) are shown. **(E)** ARPE-19 cells were infected with recombinant adenoviruses (null AD, Ad-IFITM1, and Ad-MX1) in combination. The cells were infected with cell-free VZV at an MOI of 0.01 at 48 hpi and also used for western blot analysis using antibodies to flag and GAPDH. Virus titers were determined by plaque assay at 4 dpi. Data are the averages of three independent experiments. P values are the result of Student’s t tests and error bars indicate the standard deviation from three independent experiments.

MX1-knockout ARPE-19 cell line was generated using the integrated CRISPR-Cas9 system (see Materials & Methods). MX1 was not detected in ΔMX1 ARPE-19 cells (Fig. 6A). However, other antiviral proteins CTSS, IFITM1, IRF1, IRF7, ISG15, RSAD2 were detected in ΔMX1 cells (Fig. 6A), indicating that MX1 gene was knocked out. IFN-γ-mediated anti-VZV response was assessed in the MX1-KO APRE-19 cells. When treated with IFN-γ, VZV was increased to 6×10^3^ pfu in the MX1-KO cells (Fig. 6B), demonstrating that MX1 is not the only ISGs for the inhibition of VZV replication. When MX1 was rescued by Ad-MX1 in the MX1-KO cells, IFN-γ treatment completely inhibited VZV replication (Fig. 6C). These results suggest that MX1 is a key anti-VZV restriction factor and that other ISGs also are involved in the inhibition of VZV replication.

**FIG 6.**
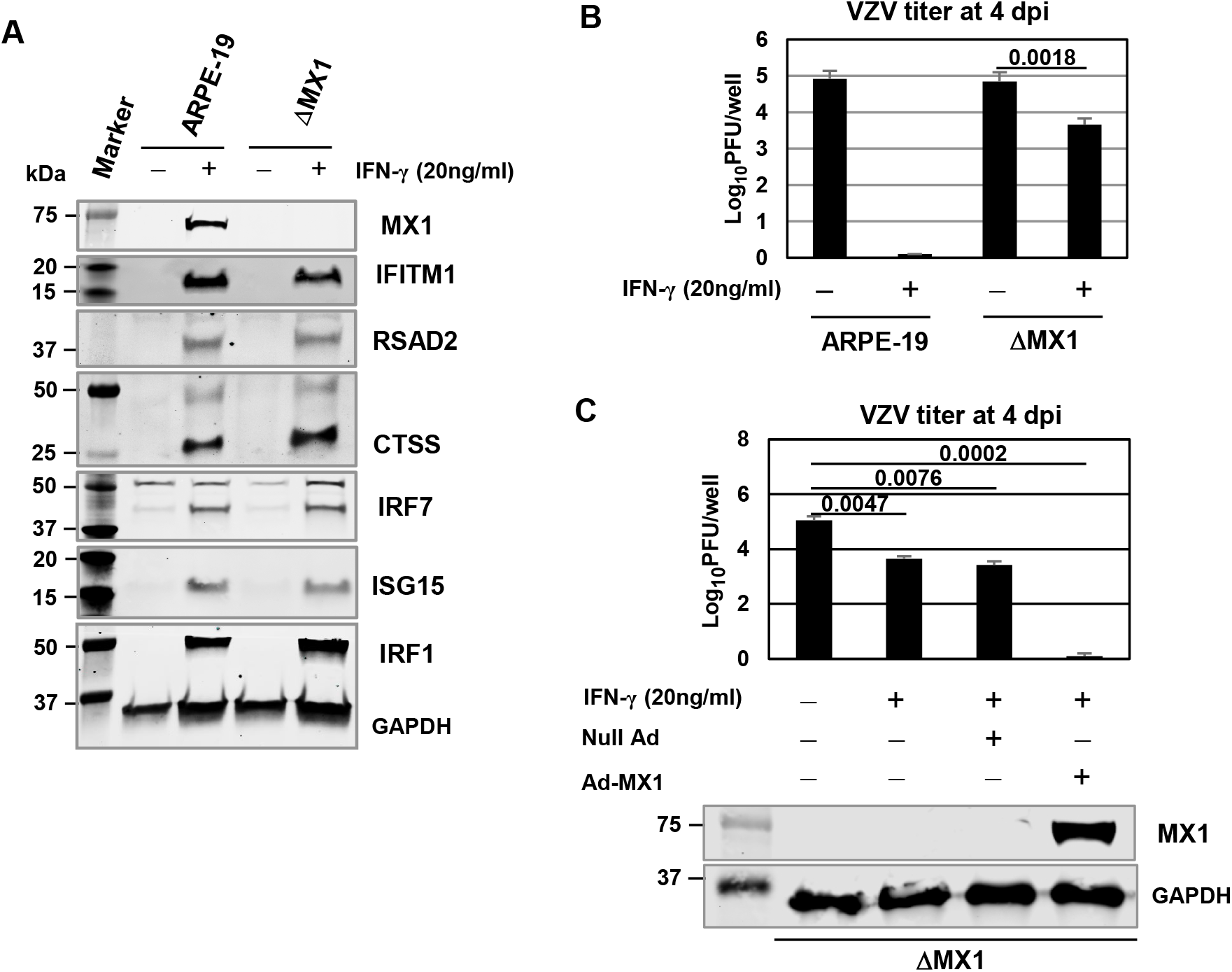
MX1 Knockout impairs IFN-γ-mediated inhibition of VZV replication in ARPE-19 cells. MX1-knockout (KO) ARPE-19 cell line was generated using the integrated CRISPR-Cas9 system (see Materials & Methods). **(A)** The MX1 proteins were not detected in the MX1-knockout cells. WT and MX1-KO ARPE-19 cells were plated in 60-mm dishes, treated with 0 (-) or 20 (+) ng/ml of IFN-γ. At 24 hpt, the cells were harvested and used for western blot analyses with antibodies. **(B)** IFN-γ-mediated anti-VZV response was assessed in MX1-KO APRE-19 cells. WT and MX1 KO ARPE-19 cells were plated in 24-well plates (3.5×10^5^), treated with 0 (-) or 20 (+) ng/ml of IFN-γ, and infected with cell-free VZV at an MOI of 0.01 at 24 hpt. Virus titers were determined by plaque assay at 4 dpi. P values are the result of Student’s t tests and error bars indicate the standard deviation from three independent experiments. **(C)** MX1 rescue was performed in MX1-KO APRE-19 cells. MX1 KO ARPE-19 cells were plated in 24-well plates (3.5×10^5^), treated with 20 (+) ng/ml of IFN-γ and infected with 30 MOI of recombinant adenoviruses (null Ad or Ad-MX1) alone or in combination. The cells were used for western blot analysis using antibodies to flag and GAPDH. Forty-eight hours later, the cells were infected with cell-free VZV at an MOI of 0.01. Virus titers were determined by plaque assay at 4 dpi. P values are the result of Student’s t tests and error bars indicate the standard deviation from three independent experiments.

### Cell-free VZV cannot infect human primary epidermal keratinocytes

To investigate whether MX1 inhibit VZV replication in human primary epidermal keratinocytes (neonatal human primary epidermal keratinocytes [HEKn] and adult human primary epidermal keratinocytes [HEKa], human primary renal mixed epithelial [REN], human primary corneal epithelial [COR], human primary bronchial epithelial [BRO]) and ARPE-19 cells were treated with 20 ng/ml of IFN-γ and the cells were infected with cell-free VZV at an MOI of 0.2 or 0.01 at 24 hpt and harvested at 4 (ARPE-19 cells) or 5 (human primary cells) dpi for western blot analyses. Two VZV immediate-early proteins IE62 and ORF63 were not detected in two human keratinocytes (HEKn and HEKa) (Fig. 7A, lane 3 and Fig. S5A). IE62 and ORF63 were detected at relatively high levels in REN (Fig. 7A, lane 6) and at very low levels in COR (Fig. 7A, lane 9) and BRO (Fig. 7A, lane 12) compared to that in ARPE-19 cells (Fig. 7A, lane 15). Insulin-degrading enzyme (IDE) is a ubiquitous protein that serves as a receptor for VZV (4, 5), indicating that receptor availability was not a limiting factor for infection across the five different human primary cells. IFN-γ treatment completely reduced the levels of the two IE proteins (Figs. 7A and S5A). Intracellular VZV was not detected in both HEKn and HEKa (Fig. S5B). We measured six ISG proteins, including MX1, in HEKn and HEKa cells in the presence or absence of IFN-γ. IFN-γ treatment strongly induced all seven ISG proteins IRF1, MX1 (human MX1), OAS2, RSAD2, GBP2, GBP4, and IFITM1 in HEKn, HEKa, and ARPE-19 cells (Fig. 7C). Interestingly, MX1 and OAS2 were expressed at high levels in the absence of IFN-γ in both HEKn and HEKa cells (Fig. 7B and 7C), indicating that basal levels of endogenous MX1 gene expression are higher in these two epidermal keratinocyte types. However, endogenous IFITM1 was expressed at very low level (Fig. 7C). Ectopic overexpression of MX1 completely inhibits VZV replication in REN cells (Fig. 7D, lane 6).

**FIG 7.**
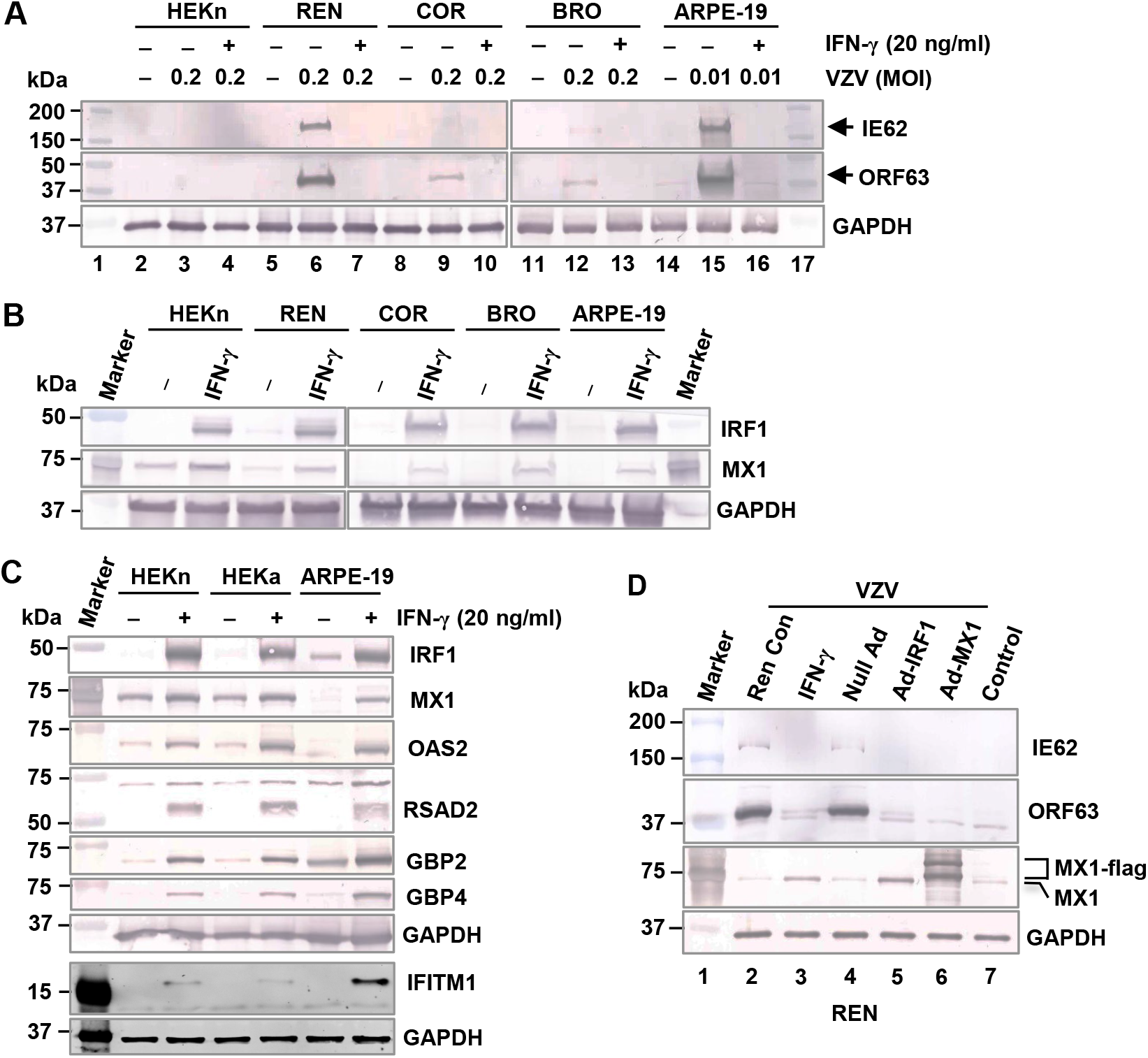
Human primary epidermal keratinocytes were more resistant to VZV infection. **(A and B)** four human primary cells (neonatal human primary epidermal keratinocytes [HEKn], human primary renal mixed epithelial [REN], human primary corneal epithelial [COR), human primary bronchial epithelial [BRO]) and ARPE-19 cells were plated in 24-well plates (3.5×10^5^), treated with 20 ng/ml of IFN-γ and the cells were infected with cell-free VZV at an MOI of 0.2 or 0.01 at 24 h posttreatment (hpt). The cells were harvested at 0 (B) or 4 dpi (A) and used for western blot analyses. **(C)** Endogenous MX1 gene expression is higher in the two epidermal keratinocytes (HEKn and adult human primary epidermal keratinocytes [HEKa]) than in ARPE-19 cells. HEKn, HEKa, and ARPE-19 cells were treated with 20 ng/ml of IFN-γ, harvested at 24 hpt, and used for western blot analyses. **(D)** Ectopic expression of MX1 completely inhibits VZV replication in primary renal mixed epithelial cells. REN cells were treated with 20 ng/ml of IFN-γ or infected with 30 MOI of recombinant adenoviruses (null Ad, Ad-IRF1, or Ad-MX1) and the cells were infected with cell-free VZV at an MOI of 0.2 at 24 hpt (or 48 hpi), harvested at 4 dpi for western blot analyses.

### Treatment with ruxolitinib or tofacitinib increases VZV replication in calcium chloride-treated primary human epidermal keratinocytes

Tofacitinib (Tofa) and Ruxolitinib (Ruxo) are inhibitors of JAK (JAK1 to 3) protein kinases and act by inhibiting cellular components of both innate and adaptive immunity [45–47]. It has been shown that treatment with tofacitinib significantly reduces gene expression of antiviral peptides such as MX1 and OAS2 in keratinocytes [61]. To examine the effect of these two JAK inhibitors on VZV replication, ARPE-19 cells were treated with Ruxo (0.7 µM) or Tofa (0.7 µM) and treated with 20 ng/ml of IFN-γ at 24 hpt. The next day, the cells were infected with cell-free VZV at an MOI of 0.01. At 4 dpi, the cells were harvested and used for western blot analyses and virus titration. In IFN-γ-treated cells, treatment with Ruxo or Tofa completely reduced the expression levels of MX1, OAS2, IRF1, and IFITM1 (Fig. 8A, lanes 5 and 7, respectively). As expected, IFN-γ treatment completely reduced viral IE62 and ORF63 gene expression (Fig. 8B, lane 4). In IFN-γ-treated cells, treatment with Tofa or Ruxo significantly increased IE62 and ORF63 expression (Fig. 8B, lanes 6 and 8, respectively) compared to those in control DMSO cells (Fig. 8B, lane 3). Very similar results were obtained for virus yields (Fig. 8C). These results indicate that the type II interferon (IFN-γ) signaling pathway was blocked by the two JAK inhibitors (tofacitinib and ruxolitinib) in VZV-infected ARPE-19 cells. Treatment with Ruxo or Tofa in HEKn cells gradually reduced the levels of MX1 and OAS2, with complete reduction achieved 5 days post-treatment (Figs. S6A and 8D). However, Ruxo or Tofa treatment failed to restore the sensitivity to VZV infection in HEKn cells (Fig. 8E).

**FIG 8.**
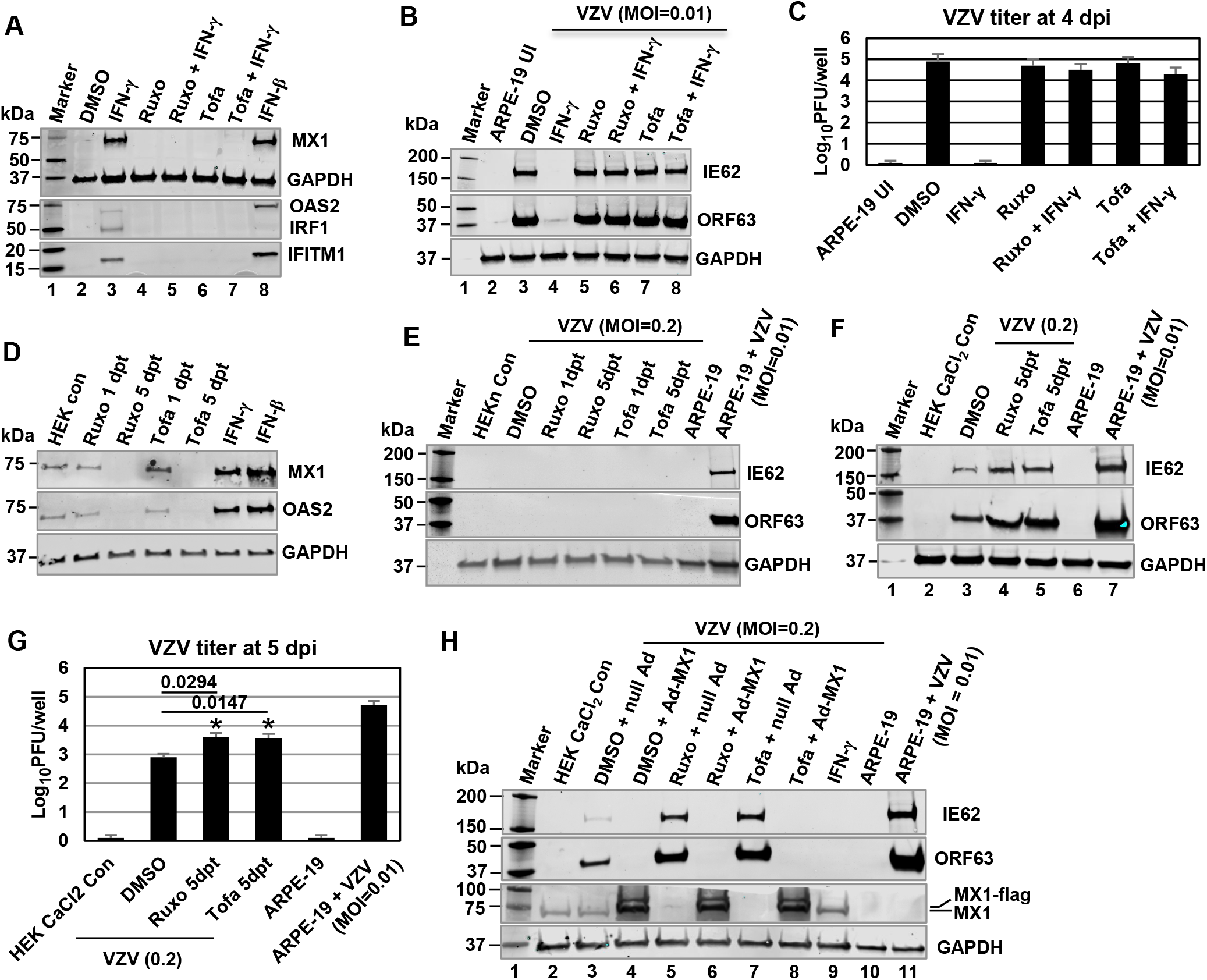
Treatment with ruxolitinib or tofacitinib increases VZV replication in calcium chloride-treated primary human epidermal keratinocytes. **(A to C)** Effect of ruxolitinib (Ruxo) and tofacitinib (Tofa) (JAK1/2 kinase inhibitors) on VZV replication in ARPE-19 cells. ARPE-19 cells were treated with Ruxo (0.7 µM; Sigma Aldrich, MO) or Tofa (0.7 µM; InvivoGen, CA) and treated with 20 ng/ml of IFN-γ at 24 hpt. The next day, the cells were infected with cell-free VZV at an MOI of 0.01. At 4 dpi, the cells were harvested and used for western blot analyses. Virus titers were determined by plaque assay at 4 dpi. UI, uninfected control. Data are the averages of three independent experiments. **(D and E)** Effect of ruxolitinib and tofacitinib (JAK1/2 kinase inhibitors) on VZV infection in neonatal human primary epidermal keratinocytes (HEKn). HEKn cells were treated with DMSO, Ruxo (0.7 µM) or Tofa (0.7 µM), infected with cell-free VZV at an MOI of 0.2 at 1 or 5 dpt. At 5 dpi, the cells were harvested and used for western blot analyses. ARPE-19 cells (3.4×10^5^) were infected with cell-free VZV at an MOI of 0.01 and harvested at 4 dpi for western blot analyses **(E, F, and H)**. **(F and G)** Effect of ruxolitinib and tofacitinib on VZV replication in HEKn cells treated with CaCl_2_. HEKn cells were plated in a 24-well plate (2.0×10^5^) in 0.6 mM CaCl_2_ containing media, treated with Ruxo (0.7 µM) or Tofa (0.7 µM) and infected with cell-free VZV (0.2 MOI) at 5 dpt. The cells were harvested at 5 dpi for western blot analysis (F). Virus titers in HEKn and ARPE-19 cells were determined by plaque assay at 5 and 4 dpi, respectively (G). P values are the result of Student’s t tests and error bars indicate the standard deviation from three independent experiments. **(H)** Ectopic overexpression of MX1 completely inhibits VZV replication in primary human epidermal keratinocytes. HEKn cells were plated in a 24-well plate (2.0×10^5^) in 0.6 mM CaCl_2_-supplemented media, treated with DMXO, Ruxo (0.7 µM) or Tofa (0.7 µM). The cells were treated with 20 ng/ml of IFN-γ or infected with 30 MOI of recombinant adenoviruses (null Ad or Ad-MX1) at 3 dpi. The cells were infected with cell-free VZV at an MOI of 0.2 at 2 dpi and harvested at 4 dpi for western blot analyses.

It has been shown that calcium regulates the differentiation of keratinocytes [48] which increased VZV gene expression, replication, and spread [49, 50]. RNA-seq analysis with calcium-induced keratinocytes showed that the differentiation status of the keratinocytes influences the replication pattern of the viral gene and protein expression [51]. VZV was able to infect HEKn treated with CaCl_2_ (Fig. 8F, lane 3 and 8G bar 2). Treatment with Ruxo or T ofa further increased VZV yields by 5.0- and 4.5-fold, respectively (Fig. 8G, bars 3 and 4, respectively). Ectopic overexpression of MX1 (human MX1) completely reduced the levels of viral IE62 and ORF63 proteins in primary human epidermal keratinocytes (Fig. 8H, lanes 4, 6, and 8). Taken together, these results demonstrate that Ruxo or Tofa treatment increases VZV replication in calcium chloride treated keratinocytes, suggesting that suppression of JAK-STAT pathways may promote uncontrolled viral replication in differentiated cells.

## DISCUSSION

Hosts have evolved numerous mechanisms to prevent primary viral infections. Interferon signaling is an important host defense mechanism against primary infection. Keratinocytes protect the human body from exogen pathogens by producing antimicrobial peptides (AMPs) and antiviral peptides but also (pro-) inflammatory chemokines and cytokines [52, 53]. Surrounding uninfected cells exhibit upregulation of IFNs, STAT1, which activates IFN-stimulated factors and other cell transactivators [54]. Interferon-stimulated proteins (ISGs) are antiviral proteins produced by cells in response to interferons that establish an “antiviral state” to block pathogen replication by targeting various stages of their life cycles. Our findings showed that basal levels of endogenous MX1 and OAS2 gene expression are higher in epidermal keratinocytes. These results suggest that keratinocytes expressing some of the antiviral genes including MX1 protect the human body from exogen pathogens.

The keratinocytes, which are the main cell type in the epidermis, are subject to cycles of proliferation, differentiation and death that allow preservation of epidermal homeostasis as well as of epidermal barrier function [55–57]. Keratinocytes retain their proliferative capacity in the basal layer; however, upon differentiation, they cease division and migrate through the suprabasal spinous and granular layers to the cornified layer. Keratinocytes are a major target of VZV replication in skin. VZV replicates, spreads among cells, and is highly cell-associated in the interfollicular basal keratinocytes but becomes infectious only when keratinocytes differentiate into the upper epidermal layers, where it accumulates as infectious cell-free virions in the cutaneous blistering lesions [51, 54, 58, 59]. It has been shown that productive VZV replication and the formation of blisters containing cell-free virions occurs only following terminal differentiation of infected basal keratinocytes [51]. The calcium-switch model of VZV infection, which allows analysis of VZV during keratinocyte differentiation also provides a useful *in vitro* model of VZV infection in human epidermis [60]. VZV gene expression was significantly increased following calcium treatment of keratinocytes [51]. The findings of this study showed that while cell-free VZV could not infect human primary epidermal keratinocytes, it was able to infect calcium chloride-differentiated keratinocytes, confirming that VZV replication is associated with cellular differentiation. Further studies are warranted to elucidate the complex molecular interactions between VZV and keratinocyte differentiation.

In human primary epidermal keratinocytes, the basal level of *MX1* gene expression is high and remains elevated following treatment with calcium chloride (Figs. 7 and S6B, respectively). Since ectopic expression of MX1 reduced VZV yields in a dose-dependent manner in ARPE-19 cells (Fig. 5E), cell-free VZV could infect calcium-stimulated keratinocytes where MX1 gene expression remained high. Tofacitinib, a Janus kinase (JAK) inhibitor, significantly reduces gene expression of antiviral peptides such as MX1 and OAS2 in keratinocytes and decreases T cell activation [61]. The suppression of MX1 expression by JAK inhibitor treatment resulted in increased VZV replication in calcium-stimulated keratinocytes (Fig. 8), indicating that endogenous MX1 influences VZV replication in keratinocytes. Because of the difficulty of cell-free VZV to infect normal keratinocytes, keratinocytes were infected by co-cultivation with VZV-infected MRC-5 cells [50] or by inoculation with cell pellets obtained via sonication of VZV-infected cells (Personal communication with Dr. Paul Kinchington, Univ. of Pittsburgh). Maintaining the keratinocytes in a low calcium media prior to VZV infection resulted in a higher initial infection [50, 51].

Human MX1 can broadly inhibit the replication of a vast range of RNA and DNA viruses [33]. Human MX1 very weakly inhibits alphaherpesvirus HSV-1 replication in mouse 3T3 cells [38]. In most hMX1-expressing cells, influenza A virus (IAV) ribonucleoprotein complexes (vRNPs) were found sequestrated together with cellular trafficking cofactors YBX1 and Rab11a in large clusters in the vicinity of the microtubule organization center (MTOC) [62]. While the anti-IAV activity of hMX1 has been intensively studied, it is not known regarding the ability of hMX1 to inhibit varicella-zoster virus (VZV). Our results showed that ectopic overexpression of MX1 or IFITM1 inhibited VZV replication both in ARPE-19 and MeWo cells. High levels of endogenous MX1 were detected in human primary epidermal keratinocytes; however, endogenous IFITM1 was expressed at a very low level (Fig. 7B and 7C). Ectopic overexpression of MX1 completely inhibits VZV gene expression both in primary renal mixed epithelial cells and in primary epidermal keratinocytes (Figs. 7 and 8, respectively). These results lead us to hypothesize that human MX1 inhibits VZV replication by sequestering viral mRNAs and proteins in the cytoplasm, a premise central to our future investigations.

Ectopic overexpression of MX1 completely inhibited VZV replication in both ARPE-19 and MeWo cells (Fig. 5). However, knockout of the MX1 gene moderately impaired IFN-γ-mediated inhibition of VZV replication in ARPE-19 cells (Fig. S6), indicating that IFN-γ treatment still moderately inhibited VZV replication in the MX1-KO ARPE-19 cell line. IFITM1 weakly inhibited VZV replication (Fig. 2), suggesting that other ISGs besides MX1 and IFITM1 are also involved in the inhibition of VZV replication. Six ISGs IRF1, CTSS, GBP4, MX1, IFIT1, and IFITM1 were selected by siRNA depletion in figure 1. Recombinant adenoviruses Ad-CTSS and Ad-IFIT1 did not reduce intracellular VZV yields (Fig. 1C). CTSS and IFIT1 siRNAs may affect expression of other genes by an off-target effect. We could not test GBP4 because a recombinant adenoviral vector expressing GBP4 was not available. In our microarray analysis, 34 ISGs were upregulated by greater than 2.5-fold at 8 h post-treatment in both ARPE-19 and MRC-5 cells compared to those of MeWo cells (Table 1). When knockdown with siGenome SMARTpool siRNAs targeting 17 ISGs (APOL2, CCL2, CMPK2, DDX58, EPST11, FBXO6, GBP1, GBP2, GBP5, IFI30, IFI35, IFI44L, NAMPT, RARRES3, TNFSF10, WARS, and XAF1; G-CUSTOM-430650 and 433679), no significant results were obtained (Fig S7; knock efficiencies were not examined). Twelve (BST2, CIITA, CLIC2, CXCL10, CXCL11, DDX60, IFI27, IFIT2, IFIT3, ISG20, RSPO3, SERPING1) of thirty-four significantly upregulated ISGs were not examined by siRNA depletion. Further work is required to select additional ISGs that are involved in the inhibition of VZV replication.

Humans possess three primary antiviral IFITMs: IFITM1, IFITM2, and IFITM3 [26]. Expression of IFITM proteins inhibits the entry of HIV-1, influenza A virus, West Nile virus, and dengue virus into cells [27, 28]. IFITM proteins reduce HIV-1 protein synthesis by specifically excluding viral transcripts from polysomes [30]. Our results showed that ectopic expression of IFITM1 reduced the protein levels of IE62, but not the level of IE62 mRNA in ARPE19 cells, suggesting that IFITM1 expression reduces the expression level of IE62 by post-transcriptional regulation. The IFITM1 protein could inhibit the translation of VZV IE62 mRNA by several mechanisms that include: (i) affecting the stability of the IE62 mRNA and (ii) the specific exclusion of IE62 mRNA from polysomes. Because IFITM1 did not reduce IE62 mRNA levels in transiently transfected or VZV-infected cells (Fig. 4), the possibility that IFITM1 affects IE62 mRNA stability is excluded. Taken together, these results suggest that IFITM1 expression reduces VZV IE62 protein synthesis by preferentially excluding viral mRNA transcripts from translation, thereby restricting viral production. These are important hypotheses for our future work.

As mentioned before, IFN-γ induce**s** the primary response gene, IRF1, which in turn activates numerous secondary response genes [23, 24]. Our findings show that IRF1 transcript levels were strongly upregulated in all three cell lines (ARPE-19, MRC-5, and MeWo) treated with IFN-γ. Our previous results showed that IFN-γ treatment increased by ∼8-fold the levels of IRF1 expression in all three cell lines [19]. However, there is no apparent defect in endogenous IRF1 activity in MeWo cells, as GBP2 and WARS were expressed at very similar levels in MeWo cells compared to ARPE-19 and MRC-5 cells (Table 1). IFN-γ and Ad-IRF1 almost completely reduced intracellular VZV yields in ARPE-19 cells, but only weakly reduced it in MeWo cells compared to control null Ad (Figs. 2 and 5), a finding consistent with our previously published results [19]. Very similar results were obtained with the same alphaherpesvirus EHV-1. Both IFN-γ and Ad-IRF1 strongly reduced EHV-1 yield in ARPE-19 cells but weakly reduced it in MeWo cells compared with the control null Ad (Fig. S4). Taken together, these results suggest that IRF1 does not properly induce antiviral ISGs in MeWo cells. Further studies are warranted to dissect what caused the impairment of IRF1-mediated ISG induction in MeWo cells.

VZV and equine herpesvirus 1 (EHV-1) are members of the subfamily *Alphaherpesvirinae* and genus *Varicellovirus* and have very similar genomic structure (group D) [63]. Our published results showed that murine IFN-γ was significantly upregulated at 8 h post-challenge in the lungs of EHV-1 pathogenic strain RacL11-challenged mice that had been immunized with attenuated EHV-1 strain KyA [64]. The administration of murine IFN-γ blocked EHV-1 replication and significantly reduced the expression levels of viral early and late regulatory proteins in the murine alveolar macrophage MH-S cells but not in murine fibroblast L-M cells [64]. Affymetrix microarray analyses of IFN-γ-treated MH-S and L-M cells revealed that five antiviral ISGs (MX1, SAMHD1, IFIT2, NAMPT, TREX1, and DDX60) were significantly upregulated only in MH-S cells [65]. It is interesting to note that MX1 gene was upregulated by 18.1-fold in MH-S cells but not in L-M cells.

In conclusion, our microarray analysis revealed that a small subset of interferon-stimulated genes (ISGs) was significantly upregulated both in ARPE-19 and MRC-5 cells compared to those in MeWo cells. The depletion of CTSS, GBP4, MX1, IFIT1, IFITM1 and IRF1 by siRNA in IFN-γ-treated cells significantly increased VZV yields. Ectopic overexpression of interferon-induced MX1 or IFITM1 inhibited VZV replication both in ARPE-19 and MeWo cells. High levels of endogenous MX1 were detected in human primary epidermal keratinocytes. Suppression of MX1 expression by JAK inhibition increased VZV replication in calcium chloride-differentiated keratinocytes. These results demonstrate that both the expression level of endogenous MX1 and the differentiation status of the keratinocytes influence VZV replication in keratinocytes. Further investigation into the functional mechanisms of IFITM1 and MX1 in VZV replication could provide important biological roles in VZV gene programming, but also a potential target for development of antiviral agents that specifically interfere with herpesvirus transcription.

## MATERIALS AND METHODS

### Viruses and cell culture

The VZV (AV92-3:L, ATCC) was cultured in ARPE-19 (CRL-2302; ATCC) cells. The KyA strain of EHV-1 was propagated in suspension cultures of L-M mouse fibroblasts [64]. Immortalized human retinal epithelial ARPE-19 cells (passage #: 24), normal human fetal lung fibroblasts MRC-5 (passage #: 10; CCL-171; ATCC), and human melanoma MeWo cells (a gift from Jeffrey Cohen, NIH) were maintained at 37°C in Dulbecco’s Modification of Eagle’s Medium with 4.5g/L glucose, L-glutamine & sodium pyruvate (10-013-CV, Corning) supplemented with 100 U/ml of penicillin, 100 µg/ml of streptomycin, nonessential amino acids, and 5% fetal bovine serum.

Neonatal human primary epidermal keratinocytes (HEKn; ATCC PCS-200-010) and adult human primary epidermal keratinocytes (HEKa; ATCC PCS-200-011) were cultured in dermal cell basal medium (ATCC, PCS-200–030) supplemented with a keratinocyte growth kit (ATCC, PCS-200–040), 100 U/ml of penicillin, and 100 µg/ml of streptomycin. Human primary renal mixed epithelial (REN; ATCC PCS-400-012), human primary corneal epithelial (COR; ATCC PCS-700-010), and human primary bronchial epithelial (BRO; ATCC PCS-300-010) were cultured in their respective basal media supplemented with growth kits.

### Preparation of cell-free VZV

Cell-free VZV was prepared by following previously published method [66, 67] with some modifications [19]. Briefly, ARPE-19 cells infected with VZV were scraped in PBS-sucrose-glutamate-serum (PSGC) buffer (5% [wt/vol] sucrose, 0.1% [wt/vol] sodium glutamate, and 10% heat-inactivated fetal calf serum in 10x PBS [68]. The cells/buffer were homogenized with a Dounce homogenizer on ice, treated with RNase A, and clarified by centrifugation at 1,500 X g for 15 min at 4°C and concentrated by using the Lenti-X concentrator (Clontech, Mountain View, CA). Cell-free VZV was aliquoted and stored in liquid nitrogen.

### Cell treatment and infection

ARPE-19, MRC-5, and MeWo cells were plated in 12-well plates (3×10^5^) or 6-well plates (7.5×10^5^), treated with recombinant human IFN-γ (CR100A, Cell Sciences, Canton, MA) at 20 ng/ml for 24 h and were incubated with 0.01 MOI of cell-free VZV for 2 h with rocking every 15 min at 37°C. After 2 h of incubation, the cells were washed with medium and cultured with fresh medium (5% FBS). At the indicated time post-infection, the cell culture was harvested for use in a plaque assay. Virus titers were determined by plaque assay [19].

### Generation of *MX1*-knockout ARPE-19 cell line using CRISPR-Cas9 system

Primers for cloning gRNAs were as follows: MX1-1 forward: caccgggagtcaatgaggtcgatgc; reverse: aaacgcatcgacctcattgactccc. MX1-2 forward: caccggatcgtgaccagatgcccgc; reverse: aaacgcgggcatctggtcacgatcc. gRNA sequences against MX1 were synthesized and cloned into lentiCRISPRv2 (Addgene #52961) according to published protocol (http://www.genome-engineering.org/gecko/) [69]. MX1 KO gRNAs were transfected into HEK-293T cells with psPAX2 (packaging plasmid; Addgene #12260) and pVSV-G (envelope plasmid; Addgene #138479) to generate lentivirus. ARPE-19 cells were transduced with serial dilutions of MX1 KO gRNA-containing lentivirus and selected with Puromycin at 1 µg/ml. MX1 KO cells were further confirmed by immunoblotting.

### Western blotting

Preparation of total extract from cells and western blot analysis were performed as previously described [70, 71], with some modifications. Lysates were mixed with 2X Laemmli sample buffer, heated to 95 °C for 3 min before running on a 4–20% Tris-Glycine gel (Bio-Rad Hercules, CA) and transferring to a NC membrane (Bio-Rad). Membranes were incubated with the indicated primary antibodies for 1 hour, incubated with secondary antibody (anti-rabbit IgG [Fc]-alkaline phosphatase [AP] conjugate) (Promega, Madison, WI) for 30 min, and visualized by incubating the membranes containing blotted protein in AP conjugate substrate (AP conjugate substrate kit, Bio-Rad) according to manufacturer’s instructions. The density of the brands on the membrane was scanned with Epson Perfection V600 Photo (Epson America, Inc., Los Alamitos, CA) and analyzed with ImageJ (Rasband, W.S., NIH, Bethesda, Maryland).

Some membranes were incubated with Secondary antibodies (goat anti-mouse [1:15,000; IRDye® 680RD, LICORbio] or goat anti-rabbit [1:15,000; IRDye® 800CW, LICORbio]) and visualized on a LI-COR Odyssey CLx.

Primary antibodies: β-actin (SC1615-R, Santa Cruz Biotech), CTSS (PA5-14271, Invitrogen), Flag pAb (Cell Signalling, Inc.), GAPDH (SC-51905, Santa Cruz Biotech., CA), GBP2 (11854-I-Ap, Proteintech, Rosemont, IL), GBP4 (17746-I-Ap, Proteintech), GFP (SC-5385, Santa Cruz Biotech.), IEP (OC33, [72]), ICP4 (58S, ATCC), IE62 pAb [73], IFITM1 (60074-I-Ap, Proteintech), IRF1 (D5E4, Cell signaling, Inc.), IRF7 (22392-I-Ap, Proteintech), ISG15 (15981-I-Ap, Proteintech), MX1 (H285, Santa Cruz Biotech., CA), OAS2 (19279-I-Ap, Proteintech), ORF63 pAb [19], RSAD2 (11833-I-Ap, Proteintech), SAMHD1 (12586-I-Ap, Proteintech),

### Luciferase reporter and mammalian expression plasmids, and recombinant adenoviruses

Plasmids were constructed and maintained in *Escherichia coli* (*E. coli*) HB101 or JM109 by standard methods [74]. Plasmids pORF61-Luc [73], pcDNAI/AMP (Invitrogen), and pCMV-IE62 [7], pSV-ICP4 [43], pSVIE [75], pICP22-SEAP (76), pC-FIT1 (pcDNA3.1 3xflag IFIT1), pC-FIT2 (pcDNA3.1 3xflag IFIT2), pC-FIT3 (pcDNA3.1 3xflag IFIT3), pC-FIT5 (pcDNA3.1 3xflag IFIT5), pC-IFITM1 (pcDNA3.1 3xflag IFITM1) (Addgene, Warertown, MA; [77]), pC-IRF1 (pCMV6-XL5, Origene Tech., Rockville, MD) have been described previously. Plasmid pICP22-SEAP has been described previously

Null Ad (ViralQuest, Inc, North Liberty, IA), Ad-GFP (CV10001, Vigene Biosciences, Rockville, MD), Ad-IRF1 (VH825936, ViGene Biosciences), Ad-MX1 (VH839486), Ad-SAMHD1 (VH893768), Ad-IFIT1 (VH83271)], Ad-GBP1 (VH860461), Ad-GBP2 (VH8070150, Ad-CTSS (VH800104), and Ad-GBP5 (VH804787), Ad-IFITM1 (VH808874, ViGene Biosciences, MD)

### Luciferase reporter assays

The luciferase reporter assay was performed by using Lipofectamine 3000 reagent (Invitrogen, San Diego, CA) according to the manufacturer’s protocol. ARPE-19 and MeWo cells were seeded at 70% confluency in 24-well plates, infected with 10 MOI of recombinant adenoviruses expressing ISGs, transfected with 0.2 pmol of expression plasmids, or 20 ng/ml of IFN-γ (Cell Sciences), and transfected with 0.07 pmol of reporter vector and 0.07 pmol of effectors in each well at 24 h post-treatment (or post-infection). Four microliters of Lipofectamine 3000 were diluted with 114 µl of Opti-MEM medium (Invitrogen). DNA and 1.5 µl of P3000 reagent were mixed with 114 µl of Opti-MEM medium. The mixed solutions were combined and incubated for 10 min at room temperature, and one-third volume was transferred into each of three wells of the cells. At 40 h post-transfection, luciferase activity was measured with a luciferase assay kit (Promega, Madison, WI) and a Polarstar Optima plate reader (BMG LABTECH Inc., Cary, NC). Reporter plasmid pICP22-SEAP was used for the internal controls. SEAP activity in culture medium was determined as previously described (78) by a chemiluminescence method using a Great Escape SEAP kit (Clontech, Palo Alto, CA).

### Small interfering RNA (siRNA) treatment

siGenome SMARTpool siRNAs targeting 11 ISGs (G-CUSTOM-430650) and nontargeting control (NTC) siRNA (Pool Cat #: D-001206-13-05) were purchased from Dharmacon (Lafayette, CO). Eleven ISGs or control siRNAs were reverse transfected into cells using Lipofectamine RNAiMAX reagent (Thermo Fisher) according to the manufacturer’s instructions. Mix 1 was prepared by adding 20 pmol of siRNA to 150 μl Opti-MEM medium (Invitrogen) and gently mixing. Mix 2 was prepared by adding 6 μl of RNAiMAX reagent to 150 μl Opti-MEM. Mixes 1 and 2 were then combined, transferred to 6-well plate, and incubated at room temperature for 5 min. ARPE-19 cells (7.5×10^5^) were then added to the well in 2.5 ml of DMEM containing 2% FBS. The cells were treated with 20 ng/ml of IFN-γ (Cell Sciences, MA) at 8h posttransfection and harvested at 24 h post-treatment for western blot analyses.

### Microarray analysis

Microarray analyses were performed as previously described [64]. Three replicates of MRC-5, ARPE-19, and MeWo cells were treated with either 0 or 20 ng/mL of human IFN-γ (Cell Sciences, MA) and harvested at 8 h post-treatment. The quality and quantity of RNA were determined, and RNA was processed with the GeneChip human Genome U133 Plus 2.0 (Affymetrix, Santa Clara, CA) at the LSUHSC-S CMTV Genomics/DNA Array Core Facility. Biotinylated cRNA was generated using the Affymetrix 3′ IVT Kit (Affymetrix, CA, USA), per the manufacturer’s instructions. Arrays were scanned using a GeneChip Scanner 3000 7G with autoloader. Pixel intensities were measured, expression signals were analyzed, and features were extracted using the commercial software package Transcriptome Analysis Console 3.0 (Affymetrix, Santa Clara, CA, USA). The data were normalized by mean values (for Spotfire pair-wise comparisons and SAM two-class comparisons) or percentile values (for GeneSpring analyses). Gene expression changes were considered significant if the p value was less than 0.05, the fold change was at least 2.0 between IFN-γ treated and untreated cells, and changes in gene expression were reproducible in all replicate comparisons. Genes expressed at different levels in untreated controls were excluded from analysis.

### Real-time RT-PCR assays

Total RNA was purified by using the RNeasy Mini kit (QIAGEN, CA) and analyzed by real-time qRT-PCR. Quantitative real-time RT-PCR (RT-qPCR) assays were performed as previously described [64]. IE62 primers (Forward, 5’-ccttggaaaccacatgatcgt-3’; Reverse, 5’-agcagaagcctcctcgacaa-3’; [79]) and GAPDH Primers (Forward, 5’-gtctcctctgacttcaacagcg-3’; Reverse, 5’-accaccctgttgctgtagccaa-3’; [79]) were synthesized from IDT (Coralville, IA). Real-time qRT-PCR amplification was carried out with the CFX96™ Real-Time PCR Detection System by using the iTaq™ Universal SYBR® Green One-Step Kit according to the manufacturer’s instructions (Bio-Rad). Each sample was assayed in triplicate.

### Microarray data accession number

The microarray data were deposited in the NCBI Gene Expression Omnibus (GEO) database under accession number GSE165112 (http://www.ncbi.nlm.nih.gov/geo).

### Statistical analysis

Data were determined by the unpaired, two tailed Student’s t-tests. Error bars represent as mean ± SD. Each experiment was carried out independently at least three times.

## SUPPLEMENTAL MATERIAL

Supplemental figures

Fig. S1 to S7.

Supplemental table

Table S1.

## ACKNOWLEDGMENTS

This research was supported by the National Institute of General Medical Sciences of the NIH under award P30GM110703. H.G. is supported by NIHR01AI189875.

We thank Drs. Paul Kinchington, University of Pittsburg and Jeffrey Cohen, NIH for providing helpful comments on cell-free VZV preparation. We thank Dr. Dennis J. O’Callaghan for helpful discussion. We also thank Dr. Stephanie E. Ander for critical reading of the manuscript and providing comments.

## Supporting information

**Fig. S1** Depletion of antiviral gene expression in ARPE-19 cells. **(A to K)** ARPE-19 cells were reverse transfected with siGenome SMARTpool siRNAs targeting 11 ISGs (G-CUSTOM-430650) or a nontargeting control (siNTC). The cells were treated with 20 ng/ml of IFN-γ (Cell Sciences, MA) at 8h posttransfection and harvested at 24 h post-treatment for western blot analyses using antibodies to IFITM1, CTSS, GBP4, IFIT1, IRF7, ISG15, OAS2, RSAD2, SAMHD1 (Proteintech, IL), MX-1 (SC50509, Santa Cruz Biotech, CA), IRF1 (D5E4, Cell Signalling Tech, Inc., MA), and β-actin (SC1615-R, Santa Cruz Biotech). The sizes of molecular mass standards (Bio-Rad, CA) are shown. The densitometry analysis was performed with ImageJ (Rasband, W.S., NIH, Bethesda, Maryland). The intensities of the ISG bands were normalized to β-actin.

**Fig. S2** ARPE-19, MRC-5, and MeWo cells were treated with 0 or 20 ng of IFN-γ and infected with 0.01 MOI of cell-free VZV at 24 h post-transfection. IE62 mRNA levels were measured *by* RT-qPCR at 3 hpi. Data are the averages of three independent experiments. *, P<0.05 for comparison with the untreated control.

**Fig. S3** IFN-γ and IRF1 reduce EHV-1 yields in a cell-line dependent manner. (**A and B**) ARPE-19 and MeWo cells were treated with IFN-γ or infected with 30 MOI of null Ad (ViralQuest, Inc., North Liberty, IA) or 30 MOI of Ad-IRF1, and infected with 0.005 MOI of EHV-1 KyA [64] at 24 h post-treatment (or 48 h post-Ad infection). Virus titers were determined at 48 hpi by plaque assay [64]. P values are the result of Student’s t tests and error bars indicate the standard deviation from three independent experiments. (**C**) ARPE-19 and MeWo cells were treated with 0 or 20 ng/ml of IFN-γ and harvested at 24 h post-treatment. ARPE-19 and MeWo cells were infected with 30 MOI of Ad-IRF1 and harvested at 48 hpi. Western blot analyses with the anti-IRF1 monoclonal antibody D5E4 (Cell Signalling Tech, Inc.) and anti-β-actin. Numbers to the left represent molecular weight standards (kDa).

**Fig. S4** MX1 strongly reduces EHV-1 yields in ARPE-19 cells. **(A and B)** ARPE-19 and MeWo cells were treated with IFN-γ or infected with 30 MOI of recombinant adenoviruses (null Ad, Ad-SAMHD1 or Ad-MX1), and infected with 0.005 MOI of EHV-1 KyA at 24 h post-treatment (or 48 h post-Ad infection). Virus titers were determined at 48 hpi by plaque assay. P values are the result of Student’s t tests and error bars indicate the standard deviation from three independent experiments.

**Fig. S5** Human primary epidermal keratinocytes cannot be infected with cell-free VZV. **(A and B)** HEKn, HEKa, and ARPE-19 cells were treated with 20 ng/ml of IFN-γ, infected with cell-free VZV (0.01 or 0.2 MOI) at 24 hpt, and harvested at 5 dpi for western blot analyses (A) and plaque assay (B). Virus titers were determined by plaque assay. Data are the averages of three independent experiments.

**Fig. S6 (A)** Treatment with ruxolitinib and tofacitinib (JAK1/2 kinase inhibitors) reduces endogenous MX1 expression in human primary epidermal keratinocytes (HEKn). HEKn cells were plated in a 24-well plate (2.0×10^5^), treated with DMSO, ruxolitinib (0.7 µM) or tofacitinib (0.7 µM). The cells were harvested at 1, 3, and 5 dpi for western blot analyses. **(B)** Treatment with ruxolitinib and tofacitinib reduces endogenous MX1 expression in primary human keratinocytes treated with CaCl_2_. HEKn cells were plated in a 24-well plate (2.0×10^5^) in 0.6 mM CaCl_2_ containing media, treated with DMSO, Ruxo (0.7 µM) or Tofa (0.7 µM). The cells were harvested at 1, 3, and 5 dpi for western blot analyses.

**Fig. S7** Depletion of APOL2, CCL2, CMPK2, DDX58, EPST11, FBXO6, GBP1, GBP2, GBP5, IFI30, IFI35, IFI44L, NAMPT, RARRES3, TNFSF10, WARS, and XAF1 in IFN-γ-treated cells does not increase VZV replication. ARPE-19 cells (3.5×10^5^, 12-well plates) were reverse transfected with siRNA targeting 17 ISGs (G-CUSTOM-430650 and 433679, Dharmacon, CO) or a nontargeting control (si-NTC, Dharmacon). At 8 hpi, the cells were treated with 20 ng/ml of IFN-γ and were infected with 0.01 MOI of VZV at 24 h post-treatment (hpt). Virus titers were determined by plaque assay at 4 dpi. Control: untreated ARPE-19 cells. P values are the result of Student’s t tests and error bars indicate the standard deviation from three independent experiments.

**Table S1. Mean fold changes of interferon-stimulated genes at 8 h post-IFN-**γ **treatment in ARPE-19, MRC-5, and MeWo cells (XLSX)**. ARPE-19, MRC-5, and MeWo cells were treated with 0 or 20 ng/mL of human IFN-γ (Cell Sciences, MA), harvested at 8 h post-treatment, and used in DNA microarray analyses using GeneChip human Genome U133 Plus 2.0 (Affymetrix, Santa Clara, CA). The mean fold change from three replicate experiments was calculated as the ratios of the average gene expression levels between the treated and untreated samples.

## REFERENCES

1. Arvin A, Gilden D. 2013. Fields Virology. 6:2015-2057. Knipe D, Howley P (ed), Fields Virology, 5th ed, Lippincott Williams & Wilkins, Philadelphia, PA

2. Cohen J. 2006. Varicella-zoster virus replication, pathogenesis, and management. Fields virology 2:2773-2818. In Knipe DM, Howley PM, Griffin DE, Lamb RA, Martin MA, Roizman B, Strauss SE (ed), Fields Viriology, 5th ed, vol 2. Lippincott Williams & Wilkins, Philadelphia, PA

3. Reichelt M, Brady J, Arvin AM. 2009.The replication cycle of varicella-zoster virus: analysis of the kinetics of viral protein expression, genome synthesis, and virion assembly at the single-cell level. J Virol 83:3904–3918.

4. Li Q, Ali MA, Cohen JI. 2006. Insulin degrading enzyme is a cellular receptor mediating varicella-zoster virus infection and cell-to-cell spread. Cell 127:305–316.

5. Kuo W, Montag AG, Rosner MR. 1993. Insulin-degrading enzyme is differentially expressed and developmentally regulated in various rat tissues. Endocrinology 132:604–611.

6. Sen N, Che X, Rajamani J, Zerboni L, Sung P, Ptacek J, Arvin AM. 2012. Signal transducer and activator of transcription 3 (STAT3) and survivin induction by varicella-zoster virus promote replication and skin pathogenesis. Proc Natl Acad Sci U S A 109:600–605.

7. Sen N, Sommer M, Che X, White K, Ruyechan WT, Arvin AM. 2010. Varicella-zoster virus immediate-early protein 62 blocks interferon regulatory factor 3 (IRF3) phosphorylation at key serine residues: a novel mechanism of IRF3 inhibition among herpesviruses. J Virol 84:9240–9253.

8. Baird NL, Bowlin JL, Hotz TJ, Cohrs RJ, Gilden D. 2015. Interferon Gamma Prolongs Survival of Varicella-Zoster Virus-Infected Human Neurons In Vitro. J Virol 89(14):7425–7.

9. Feduchi E, Alonso MA, Carrasco L. 1989. Human gamma interferon and tumor necrosis factor exert a synergistic blockade on the replication of herpes simplex virus. J Virol 63:1354–9.

10. Cantin E, Tanamachi B, Openshaw H. 1999. Role for gamma interferon in control of herpes simplex virus type 1 reactivation. J Virol 73:3418–23.

11. Cantin EM, Hinton DR, Chen J, Openshaw H. 1995. Gamma interferon expression during acute and latent nervous system infection by herpes simplex virus type 1. J Virol 69:4898–905.

12. Burke JD, Young HA. 2019. IFN-γ: a cytokine at the right time, is in the right place. Semin Immunol 43:1–8.

13. Jorgovanovic D, Song M, Wang L, Zhang Y. 2020. Roles of IFN-γ in tumor progression and regression: a review. Biomark Res Sep 29;8:49.

14. van Boxel-Dezaire AH, Stark GR. 2007. Cell type-specific signaling in response to interferon-gamma. Curr Top Microbiol Immunol 316:119–54.

15. Ivashkiv LB. 2018. IFN-γ: signalling, epigenetics and roles in immunity, metabolism, disease and cancer immunotherapy. Nat Rev Immunol 18(9):545–558.

16. Jenkins DE, Redman RL, Lam EM, Liu C, Lin I, Arvin AM. 1998. Interleukin (IL)-10, IL-12, and interferon-γ production in primary and memory immune responses to varicella-zoster virus. J Infect Dis 178(4):940–8.

17. Torigoe S, Ihara T, Kamiya H. 2000. IL-12, IFN-γ, and TNF-α released from mononuclear cells inhibit the spread of varicella-zoster virus at an early stage of varicella. Microbiol Immunol 44(12):1027–31.

18. Sen N, Sung P, Panda A, Arvin AM. 2018. Distinctive roles for type I and type II interferons and interferon regulatory factors in the host cell defense against varicella-zoster virus. J Virol 92:e01151–18.

19. Shakya AK, O’Callaghan DJ, Kim SK. 2019. Interferon gamma inhibits varicella-zoster virus replication in a cell line-dependent manner. J Virol 93(12):e00257–19.

20. Levin MJ, Smith JG, Kaufhold RM, Barber D, Hayward AR, Chan CY, Chan IS, Li DJ, Wang W, Keller PM, Shaw A, Silber JL, Schlienger K, Chalikonda I, Vessey SJ, Caulfield MJ. 2003. Decline in varicella-zoster virus (VZV)-specific cell-mediated immunity with increasing age and boosting with a high-dose VZV vaccine. J Infect Dis 188(9):1336–44.

21. Schroder K, Hertzog PJ, Ravasi T, Hume DA. 2004. Interferon-gamma: an overview of signals, mechanisms and functions. J Leukoc Biol 75:163–89.

22. Bach EA, Aguet M, Schreiber RD. 1997. The IFN gamma receptor: a paradigm for cytokine receptor signaling. Annu Rev Immunol 15:563–91.

23. Schoggins JW, Wilson SJ, Panis M, Murphy MY, Jones CT, Bieniasz P, Rice CM. 2011. A diverse range of gene products are effectors of the type I interferon antiviral response. Nature 472:481–5. Erratum in: Nature. 2015;525(7567):144.

24. Boehm U, Klamp T, Groot M, Howard J. 1997. Cellular responses to interferon-γ. Ann Rev Immunol 15:749–795.

25. Fensterl V, Sen GC. 2015. Interferon-induced Ifit proteins: their role in viral pathogenesis. J Virol 89:2462–2468.

26. Bailey CC, Zhong G, Huang IC, Farzan M. 2014. IFITM-Family Proteins: The Cell’s First Line of Antiviral Defense. Annu Rev Virol 1:261–283.

27. Brass AL, Huang IC, Benita Y, John SP, Krishnan MN, Feeley EM, Ryan BJ, Weyer JL, van der Weyden L, Fikrig E, Adams DJ, Xavier RJ, Farzan M, Elledge SJ. 2009. The IFITM proteins mediate cellular resistance to influenza A H1N1 virus, West Nile virus, and dengue virus. Cell 139:1243–1254.

28. Lu J, Pan Q, Rong L, He W, Liu SL, Liang C. 2011. The IFITM proteins inhibit HIV-1 infection. J Virol 85:2126–2137. Erratum in: J Virol. 2011;85(8):4043. He, Wei [added].

29. Amini-Bavil-Olyaee S, Choi YJ, Lee JH, Shi M, Huang IC, Farzan M, Jung JU. 2013. The Antiviral Effector IFITM3 Disrupts Intracellular Cholesterol Homeostasis to Block Viral Entry. Cell Host Microbe 13(4):452–64. Erratum in: Cell Host Microbe. 2013;14(5):600-1.

30. Lee WYJ, Fu RM, Chen L, Sloan RD. 2018. The IFITM proteins inhibit HIV-1 protein synthesis. Scientific Reports 8:14551.

31. Haller O, Kochs G. 2002. Interferon-induced mx proteins: dynamin-like GTPases with antiviral activity. Traffic 3:710 –717.

32. Haller O, Staeheli P, Schwemmle M, Kochs G. 2015. Mx GTPases: dynamin-like antiviral machines of innate immunity. Trends Microbiol 23:154–163.

33. Verhelst J, Hulpiau P, Saelens X. 2013. Mx proteins: antiviral gatekeepers that restrain the uninvited. Microbiol Mol Biol Rev 77:551–566. Erratum in: Microbiol Mol Biol Rev. 2014;78(1):198.

34. Schwab LSU, Villalón-Letelier F, Tessema MB, Londrigan SL, Brooks AG, Hurt A, Coch C, Zillinger T, Hartmann G, Reading PC. 2022. Expression of a Functional Mx1 Protein Is Essential for the Ability of RIG-I Agonist Prophylaxis to Provide Potent and Long-Lasting Protection in a Mouse Model of Influenza A Virus Infection. Viruses 14(7):1547.

35. Tessema MB, Farrukee R, Andoniou CE, Degli-Esposti MA, Oates CV, Barnes JB, Wakim LM, Brooks AG, Londrigan SL, Reading PC. 2022. Mouse Mx1 Inhibits Herpes Simplex Virus Type 1 Genomic Replication and Late Gene Expression *In Vitro* and Prevents Lesion Formation in the Mouse Zosteriform Model. J Virol 96(12):e0041922.

36. Kochs G, Janzen C, Hohenberg H, Haller O. 2002. Antivirally active MxA protein sequesters La Crosse virus nucleocapsid protein into perinuclear complexes. Proc Natl Acad Sci U S A 99:3153–3158.

37. Frese M, Kochs G, Feldmann H, Hertkorn C, Haller O. 1996. Inhibition of bunyaviruses, phleboviruses, and hantaviruses by human MxA protein. J Virol Feb;70(2):915–23.

38. Ku CC, Che XB, Reichelt M, Rajamani J, Schaap-Nutt A, Huang KJ, Sommer MH, Chen YS, Chen YY, Arvin AM. 2011. Herpes simplex virus-1 induces expression of a novel MxA isoform that enhances viral replication. Immunol Cell Biol 89:173–182.

39. Diamond MS, Farzan M. 2013. The broad-spectrum antiviral functions of IFIT and IFITM proteins. Nat Rev Immunol 13 (1): 46–57.

40. Grundy FJ, Baumann RP, O’Callaghan DJ. 1989. DNA sequence and comparative analyses of the equine herpesvirus type 1 immediate early gene. Virology 172: 223–236.

41. Schwyzer M, Vlček Č, Menekse O, Fraefel C, Pačes V. 1993. Promoter, spliced leader, and coding sequence for BICP4, the largest of the immediate-early proteins of bovine herpesvirus 1. Virology 197:349–357.

42. Wu CL, Wilcox KW. 1991. The conserved DNA-binding domains encoded by the herpes simplex virus type 1 ICP4, pseudorabies virus IE180, and varicella-zoster virus ORF62 genes recognize similar sites in the corresponding promoters. J Virol 65:1149–1159.

43. Kim SK, Shakya AK, O’Callaghan DJ. 2016. Full *trans*-activation mediated by the immediate-early protein of equine herpesvirus 1 requires a consensus TATA box, but not its cognate binding sequence. Virus Res 211:222–232.

44. Smith R, Caughman G, O’Callaghan DJ. 1992. Characterization of the regulatory functions of the equine herpesvirus 1 immediate-early gene product. J Virol 66:936–945.

45. Strand V, Kavanaugh A, Kivitz AJ, van der Heijde D, Kwok K, Akylbekova E, Soonasra A, Snyder M, Connell C, Bananis E, Smolen JS. 2018. Long-term radiographic and patient-reported outcomes in patients with rheumatoid arthritis treated with tofacitinib: ORAL start and ORAL scan post-hoc analyses. Rheumatol Ther 5(2):341–353.

46. Lamba M, Wang R, Fletcher T, Alvey C, Kushner J, Stock TC. 2016. Extendedrelease once-daily formulation of tofacitinib: evaluation of pharmacokinetics compared with immediate-release tofacitinib and impact of food. J Clin Pharmacol 56(11):1362–1371.

47. Greenfield G, McPherson S, Mills K, McMullin MF. 2018. The ruxolitinib effect: understanding how molecular pathogenesis and epigenetic dysregulation impact therapeutic efficacy in myeloproliferative neoplasms. J Transl Med 16(1):360.

48. Bikle DD, Ng D, Tu CL, Oda Y, Xie Z. 2001. Calcium- and vitamin D-regulated keratinocyte differentiation. Mol Cell Endocrinol 177(1-2):161–71.

49. Tommasi C, Breuer J. 2022. The Biology of Varicella-Zoster Virus Replication in the Skin. Viruses 14(5):982.

50. Sexton CJ, Navsaria HA, Leigh IM, Powell K. 1992. Replication of varicella zoster virus in primary human keratinocytes. J Med Virol 38:260–264.

51. Jones M, Dry IR, 2014. RNA-seq Analysis of Host and Viral Gene Expression Highlights Interaction between Varicella Zoster Virus and Keratinocyte Differentiation. PLoS Pathog. 2014;10:e1003896. Erratum in: PLoS Pathog 10(7):e1004313.

52. Taniguchi K, Arima K, Masuoka M, Ohta S, Shiraishi H, Ontsuka K, Suzuki S, Inamitsu M, Yamamoto KI, Simmons O, Toda S, Conway SJ, Hamasaki Y, Izuhara K. 2014. Periostin controls keratinocyte proliferation and differentiation by interacting with the paracrine IL-1alpha/IL-6 loop. J Invest Dermatol 134(5):1295–1304.

53. Peters JH, Tjabringa GS, Fasse E, de Oliveira VL, Schalkwijk J, Koenen HJ, Joosten I. 2013. Co-culture of healthy human keratinocytes and T-cells promotes keratinocyte chemokine production and RORgammat-positive IL-17 producing T-cell populations. J Dermatol Sci 69(1):44–53.

54. Zerboni L, Sen N, Oliver SL, Arvin AM. 2014. Molecular mechanisms of varicella zoster virus pathogenesis. Nat Rev Microbiol 12(3):197–210.

55. Candi E, Schmidt R, Melino G. 2005. The cornified envelope: A model of cell death in the skin. Nat Rev Mol Cell Biol 6(4):328–40.

56. Blanpain C, Fuchs E. 2009. Epidermal homeostasis: A balancing act of stem cells in the skin. Nat Rev Mol Cell Biol 10:207–217.

57. Proksch E, Brandner JM, Jensen JM. 2008. The skin: An indispensable barrier. Exp Dermatol 17:1063–1072.

58. Gershon MD, Gershon AA. 2010. VZV infection of keratinocytes: Production of cell-free infectious virions in vivo. Curr Top Microbiol Immunol 342:173–88.

59. Chen JJ, Zhu Z, Gershon AA, Gershon MD. 2004. Mannose 6-phosphate receptor dependence of varicella zoster virus infection in vitro and in the epidermis during varicella and zoster. Cell 119(7):915–26.

60. Tommasi C, Rogerson C, Depledge DP, Jones M, Naeem AS, Venturini C, Frampton D, Tutill HJ, Way B, Breuer J, O’Shaughnessy RFL. 2020. Kallikrein-Mediated Cytokeratin 10 Degradation Is Required for Varicella Zoster Virus Propagation in Skin. J Invest Dermatol 140(4):774–784.e11.

61. Hawerkamp HC, Domdey A, Radau L, Sewerin P, Oláh P, Homey B, Meller S. 2021. Tofacitinib downregulates antiviral immune defence in keratinocytes and reduces T cell activation. Arthritis Res Ther 23(1):144.

62. McKellar J, Cadènes J, García de Gracia F, Aubé C, Chaves Valadão AL, Tauziet M, Arnaud-Arnould M, Rebendenne A, Kociánová L, Labaronne E, Ricci EP, Delaval B, Gaudin R, Naffakh N, Gallois-Montbrun S, Moncorgé O, Goujon C. 2025. Human MX1 orchestrates the cytoplasmic sequestration of neosynthesized influenza A virus vRNPs. Proc Natl Acad Sci U S A 122(41):e2418935122.

63. Roizman B, Pellet PE. 2001. The Family of Herpesviridae: a brief introduction., p. 2381–2397. In D. M. Knipe and P. M. Howley (ed.), Field Virology, Fourth ed. Lippincott Williams and Wilkins., Philadelphia, PA

64. Kim SK, Shakya AK, O’Callaghan DJ. 2016. Immunization with attenuated equine herpesvirus 1 strain KyA induces innate immune responses that protect mice from lethal challenge. J Virol 90:8090–8104.

65. Kim SK. Shakya AK, O’Callaghan DJ. 2021. Interferon gamma inhibits equine herpesvirus 1 replication in a cell line-dependent Manner. Pathogens 10(4):484.

66. Sloutskin A, Goldstein RS. 2014. Laboratory preparation of Varicella-Zoster Virus: concentration of virus-containing supernatant, use of a debris fraction and magnetofection for consistent cell-free VZV infections. J Virol Methods 206:128–32.

67. Sloutskin A, Kinchington PR, Goldstein RS. 2013. Productive vs non-productive infection by cell-free varicella zoster virus of human neurons derived from embryonic stem cells is dependent upon infectious viral dose. Virology 443:285–93.

68. Harper DR, Mathieu N, Mullarkey J. 1998. High-titre, cryostable cell-free varicella zoster virus. Arch Virol 143(6):1163–70.

69. Lane RK, Guo H., Fisher AD, Diep J, Lai Z, Chen Y, Upton JW, Carette J, Mocarski ES, Kaiser WJ. 2020. Necroptosis-based CRISPR knockout screen reveals Neuropilin-1 as a critical host factor for early stages of murine cytomegalovirus infection. Proc Natl Acad Sci U S A 117(33):20109–20116.

70. Kim SK, Buczynski KA, Caughman GB, O’Callaghan DJ. 2001. The equine herpesvirus 1 immediate-early protein interacts with EAP, a nucleolar-ribosomal protein. Virology 279:173–184.

71. Kim SK, Ahn BC, Albrecht RA, O’callaghan DJ. 2006. The unique IR2 protein of equine herpesvirus 1 negatively regulates viral gene expression. J Virol 80:5041–5049.

72. Harty RN, O’Callaghan DJ. 1991. An early gene maps within and is 3’coterminal with the immediate-early gene of equine herpesvirus 1. J Virol 65:3829–3838.

73. Kim SK, Shakya AK, Kim S, O’Callaghan DJ. 2016. Functional characterization of the serine-rich tract of varicella-zoster virus IE62. J Virol 90:959–971.

74. Sambrook J, Fritsh E, Maniatis T. 1989. Molecular cloning: A laboratory manual 2nd ed Cold Spring Harbor Press New York

75. Smith R, Caughman G, O’Callaghan D. 1992. Characterization of the regulatory functions of the equine herpesvirus 1 immediate-early gene product. J Virol 66:936–945.

76. Bedadala GR, Pinnoji RC, Hsia S-CV. 2007. Early growth response gene 1 (Egr-1) regulates HSV-1 ICP4 and ICP22 gene expression. Cell Research 17:546–555.

77. Katibah, GE, Lee HJ, Huizar JP, Vogan JM, Alber T, Collins K. 2013. tRNA binding, structure, and localization of the human interferon-induced protein IFIT5. Mol Cell 49(4):743–750.

78. Kim SK, Kim S, Dai G, Zhang Y, Ahn BC, O’Callaghan DJ. 2011. Identification of functional domains of the IR2 protein of equine herpesvirus 1 required for inhibition of viral gene expression and replication. Virology 417:430–442.

79. Depledge DP, Ouwendijk WJD, Sadaoka T, Braspenning SE, Mori Y, Cohrs RJ, Verjans G MGM, Breuer JA. 2018. spliced latency-associated VZV transcript maps antisense to the viral transactivator gene 61. Nat Commun 9:1167.

