## Supplementary material for "Interferon gamma-induced MX1 and IFITM1 inhibit Varicella-Zoster Virus Replication": Supplemetal Figures: VZV_Suppl Figures_Kim-Bio.pdf

#### Fig. S1

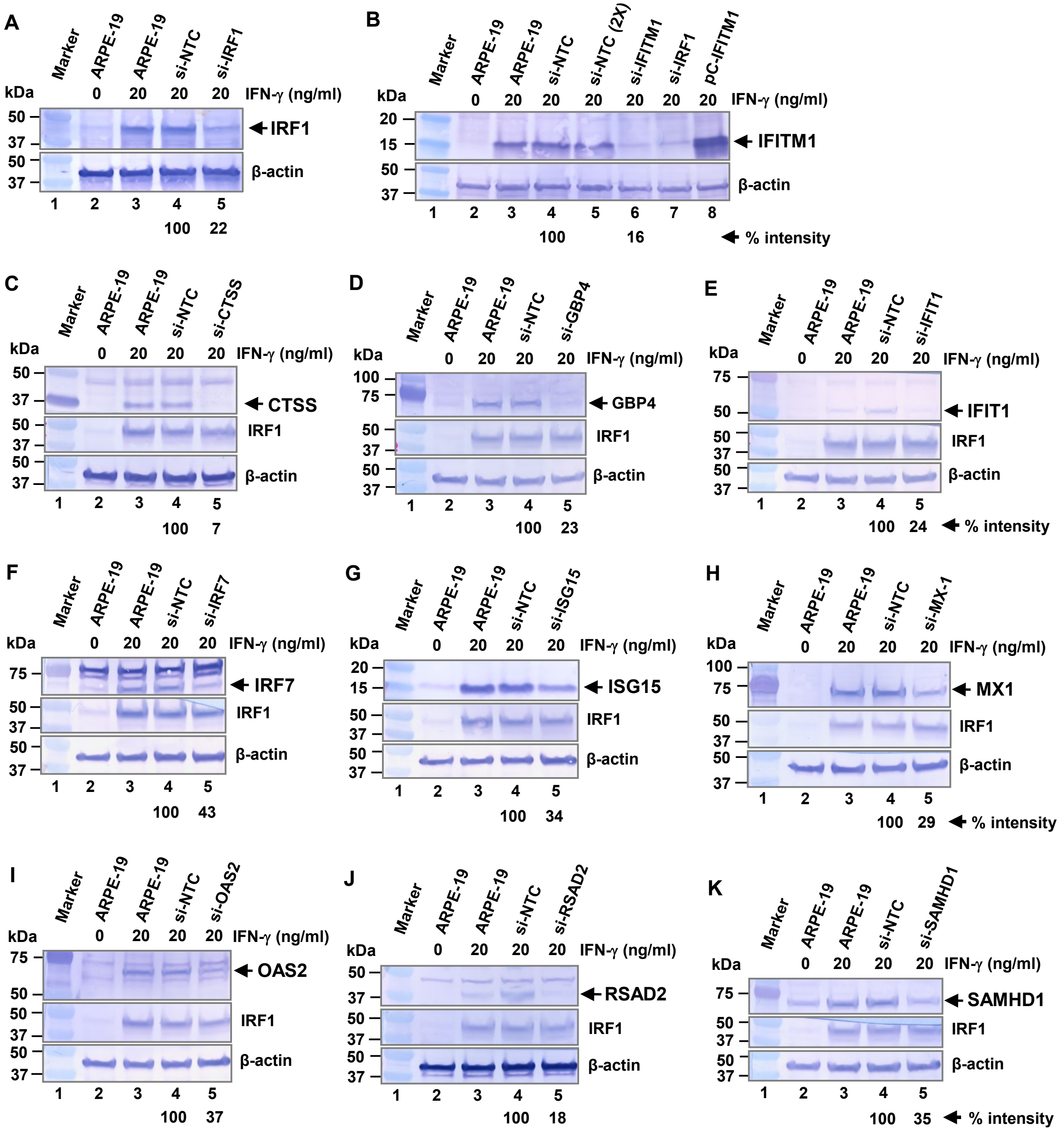

A
Marker ARPE-19 ARPE-19 si-NTC si-IRF1
kDa 0 20 20 20 IFN-γ (ng/ml)
50 37 IRF1
50 37 β-actin
1 2 3 4 5
100 22
B
Marker ARPE-19 ARPE-19 si-NTC si-NTC (2X) si-IFITM1 si-IRF1 pC-IFITM1
kDa 0 20 20 20 20 20 20 IFN-γ (ng/ml)
20 15 IFITM1
50 37 β-actin
1 2 3 4 5 6 7 8
100 16 % intensity
C
Marker ARPE-19 ARPE-19 si-NTC si-CTSS
kDa 0 20 20 20 IFN-γ (ng/ml)
50 37 CTSS
50 37 IRF1
50 37 β-actin
1 2 3 4 5
100 7
D
Marker ARPE-19 ARPE-19 si-NTC si-GBP4
kDa 0 20 20 20 IFN-γ (ng/ml)
100 75 GBP4
50 37 IRF1
50 37 β-actin
1 2 3 4 5
100 23
E
Marker ARPE-19 ARPE-19 si-NTC si-IFIT1
kDa 0 20 20 20 IFN-γ (ng/ml)
75 50 IFIT1
50 37 IRF1
50 37 β-actin
1 2 3 4 5
100 24 % intensity
F
Marker ARPE-19 ARPE-19 si-NTC si-IRF7
kDa 0 20 20 20 IFN-γ (ng/ml)
75 IRF7
50 37 IRF1
50 37 β-actin
1 2 3 4 5
100 43
G
Marker ARPE-19 ARPE-19 si-NTC si-ISG15
kDa 0 20 20 20 IFN-γ (ng/ml)
20 15 ISG15
50 37 IRF1
50 37 β-actin
1 2 3 4 5
100 34
H
Marker ARPE-19 ARPE-19 si-NTC si-MX-1
kDa 0 20 20 20 IFN-γ (ng/ml)
100 75 MX1
50 37 IRF1
50 37 β-actin
1 2 3 4 5
100 29 % intensity
I
Marker ARPE-19 ARPE-19 si-NTC si-OAS2
kDa 0 20 20 20 IFN-γ (ng/ml)
75 50 OAS2
50 37 IRF1
50 37 β-actin
1 2 3 4 5
100 37
J
Marker ARPE-19 ARPE-19 si-NTC si-RSAD2
kDa 0 20 20 20 IFN-γ (ng/ml)
50 37 RSAD2
50 37 IRF1
50 37 β-actin
1 2 3 4 5
100 18
K
Marker ARPE-19 ARPE-19 si-NTC si-SAMHD1
kDa 0 20 20 20 IFN-γ (ng/ml)
75 SAMHD1
50 37 IRF1
50 37 β-actin
1 2 3 4 5
100 35 % intensity

### Fig. S2

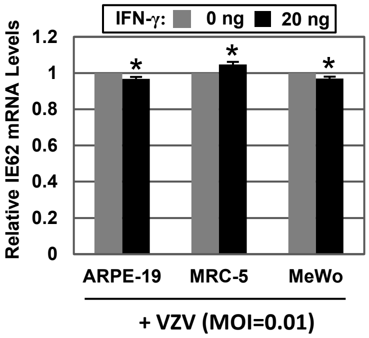
IFN-γ: 0 ng 20 ng
Relative IE62 mRNA Levels
1.2
1
0.8
0.6
0.4
0.2
0
*
*
*
ARPE-19
MRC-5
MeWo
+ VZV (MOI=0.01)

#### Fig. S3

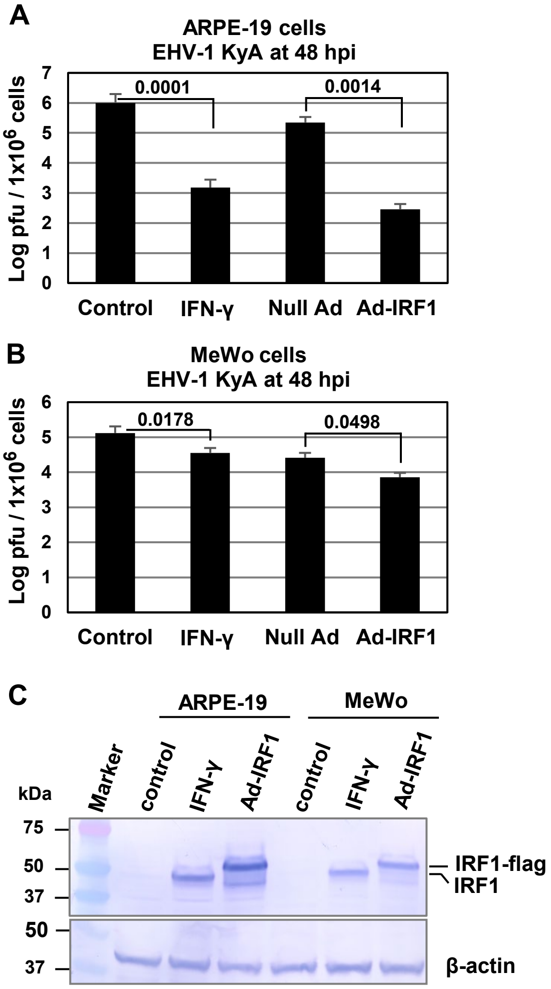
A
ARPE-19 cells
EHV-1 KyA at 48 hpi
Log pfu / 1x10^6 cells
0.0001
0.0014
Control
IFN-γ
Null Ad
Ad-IRF1
B
MeWo cells
EHV-1 KyA at 48 hpi
Log pfu / 1x10^6 cells
0.0178
0.0498
Control
IFN-γ
Null Ad
Ad-IRF1
C
ARPE-19
MeWo
kDa
Marker
control
IFN-γ
Ad-IRF1
control
IFN-γ
Ad-IRF1
75
50
37
50
37
IRF1-flag
IRF1
β-actin

### Fig. S4

A

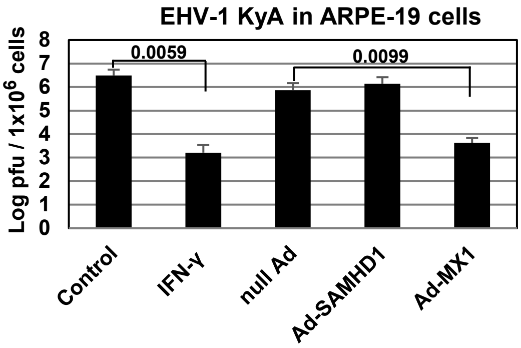
EHV-1 KyA in ARPE-19 cells
Log pfu / 1x10^6 cells
0.0059
0.0099
Control
IFN-γ
null Ad
Ad-SAMHD1
Ad-MX1

B

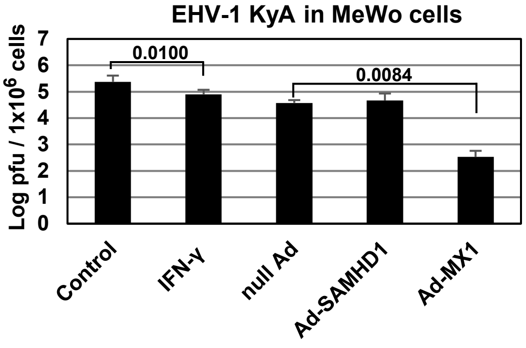
EHV-1 KyA in MeWo cells
Log pfu / 1x10^6 cells
0.0100
0.0084
Control
IFN-γ
null Ad
Ad-SAMHD1
Ad-MX1

### Fig. S5

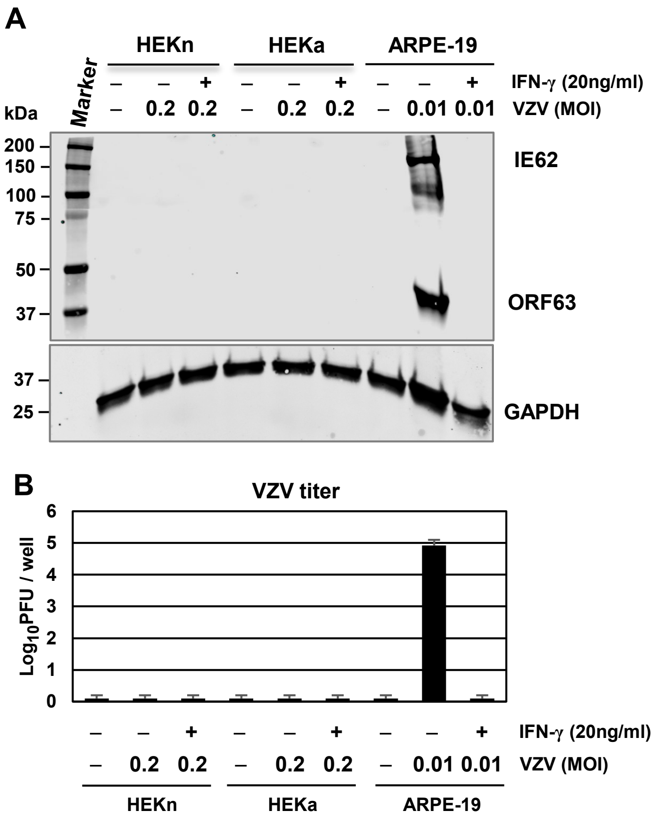
A
HEKn
HEKa
ARPE-19
Marker
IFN-γ (20ng/ml)
VZV (MOI)
kDa
200
150
100
75
50
37
IE62
ORF63
37
25
GAPDH
B
VZV titer
Log10PFU / well
6
5
4
3
2
1
0
– – + – – + – – +
– 0.2 0.2 – 0.2 0.2 – 0.01 0.01
IFN-γ (20ng/ml)
VZV (MOI)
HEKn
HEKa
ARPE-19

### Fig. S6

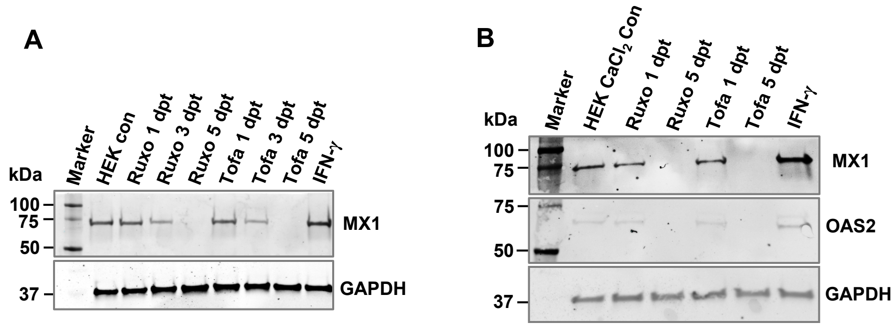

A
kDa
Marker
HEK con
Ruxo 1 dpt
Ruxo 3 dpt
Ruxo 5 dpt
Tofa 1 dpt
Tofa 3 dpt
Tofa 5 dpt
IFN-γ
100
75
50
37
MX1
GAPDH
B
kDa
Marker
HEK CaCl2 Con
Ruxo 1 dpt
Ruxo 5 dpt
Tofa 1 dpt
Tofa 5 dpt
IFN-γ
100
75
75
50
37
MX1
OAS2
GAPDH

### Fig. S7

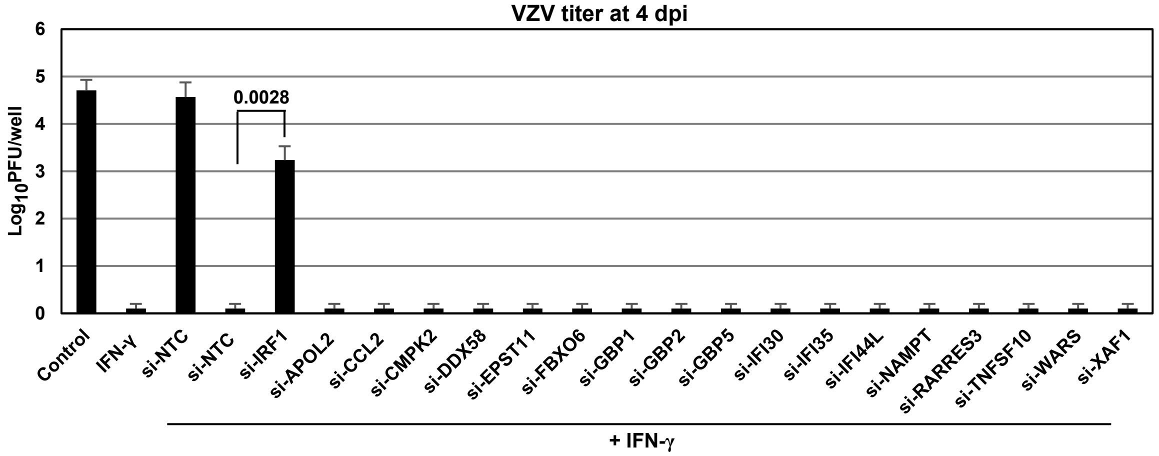

VZV titer at 4 dpi
Log10PFU/well
6
5
4
3
2
1
0
0.0028
Control
IFN-γ
si-NTC
si-NTC
si-IRF1
si-APOL2
si-CCL2
si-CMPK2
si-DDX58
si-EPST11
si-FBXO6
si-GBP1
si-GBP2
si-GBP5
si-IFI30
si-IFI35
si-IFI44L
si-NAMPT
si-RARRES3
si-TNFSF10
si-WARS
si-XAF1
+ IFN-γ
